# A macro-ecological approach to predators’ functional response

**DOI:** 10.1101/832220

**Authors:** Matthieu Barbier, Laurie Wojcik, Michel Loreau

## Abstract

Predation often deviates from the law of mass action: many micro- and meso-scale experiments have shown that consumption saturates with resource abundance, and decreases due to interference between consumers. But does this observation hold at macro-ecological scales, spanning many species and orders of magnitude in biomass? If so, what are its consequences for large-scale ecological patterns and dynamics?

We perform a meta-analysis of predator-prey pairs of mammals, birds and reptiles, and show that predation losses appear to increase, not as the product of predator and prey densities following the Lotka-Volterra (mass action) model, but rather as the square root of that product. This suggests a phenomenological power-law expression of the effective cross-ecosystem functional response. We discuss whether the same power-law may hold dynamically within an ecosystem, and assuming that it does, we explore its consequences in a simple food chain model. The empirical exponents fall close to the boundary between regimes of donor and consumer limitation. Exponents on this boundary are singular in multiple ways. First, they maximize predator abundance and some stability metrics. Second, they create proportionality relations between biomass and productivity, both within and between trophic levels. These intuitive relations do not hold in general in mass action models, yet they are widely observed empirically. These results provide evidence of mechanisms limiting predation across multiple ecological scales. Some of this evidence was previously associated with donor control, but we show that it supports a wider range of possibilities, including forms of consumer control. As limiting consumption counter-intuitively allows larger populations, it is worthwhile to reconsider whether the observed functional response arises from microscopic mechanisms, or could hint at selective pressure at the population level.

This article has been peer-reviewed and recommended by Peer Community In Ecology (DOI: 10.24072/pci.ecology.100051)

## 1 Introduction

Many dynamical food web models attempt to represent population-level dynamics by zooming out from the microscopic complexity of individual predator and prey behavior, physiology and ecology. Since Lindeman [38], these models have shared the same fundamental structure, i.e. a balance of energy gains and losses from predation and from non-trophic processes. The hidden complexity and specificity of a given ecosystem is generally summarized in a simple function, the predator functional response, which describes how fluxes between trophic levels (predation) depend on stocks at each level (predator and prey abundances). The most basic assumption is *mass action*: encounter probability increases as the product of predator and prey density. But natural settings often deviate from this baseline [32]. The expression of the functional response has been the topic of extensive empirical and theoretical inquiries, and even major debates [5, 3]. When we aim at large-scale prediction, such as a management model for the regional populations of predator and prey species, the functional response in our equations does not represent the concrete process of predation – instead, its purpose is to express the cumulative effect of many predation events on the dynamics of these aggregate variables. For instance, we may hope to predict whether an entire regional population will show a strong response to invasions or exploitation, such as a trophic cascade or collapse [43]. Yet, we often simply resort to functional forms observed at a microscopic scale, such as the saturating functional response of a single predator, and refit their parameters at the scale relevant to our prediction.

But there is no reason to expect that a single form of functional response, whether mechanistic or phenomenological, will be appropriate to model all ecological processes [4, 11]. We may have to derive different expressions for different questions and scales, from short-term laboratory experiments [57], through multi-generation predator-prey cycles [55], to macro-scale food web properties such as biomass pyramids and stability [9]. Here we propose a new, empirically-motivated and phenomenological formulation of the functional response at large spatio-temporal scales. By contrast with mechanistic theories [22, 44, 23, 48], our aim is not to ascertain the most realistic expression of predation rates, but to find a simple expression that can mimick observed rates across multiple scales, and then to understand its consequences for other food chain properties. We first perform a meta-analysis of predation rates observed for higher vertebrates in the field, covering a range of species, study durations and areas. In the existing literature, most measurements of the functional response (predation rate as a function of predator and prey density) come from feeding experiments, restricted to time scales far shorter than the predator’s generation time [53, 44]. While these experimental measurements are valuable for mechanistic models of specific populations, they may be misleading when extrapolating to macro-scale dynamics [4]. For the latter, we need a phenomenological expression that exhibits some form of scale invariance, so that it may hold across different levels of aggregation. We propose a power-law expression similar to the Cobb-Douglas production function in economics [18]. We then find best-fit exponents suggesting a square root dependence of predation on both predator and prey density. These exponents are starkly lower than those expected from the simple mass action (Lotka-Volterra) model, suggesting that hidden mechanisms strongly limit predation.

This empirical trend can only be interpreted as a meaningful functional response if it can be used to predict large-scale dynamics. This raises multiple questions. Knowing that local populations undergo many complex processes on short time scales, is it possible to write an effective model that approximates the dynamics of abundances aggregated over years and over whole landscapes? What kind of equations and nonlinearities should be used in this effective model? These questions have been raised in prior studies, mainly within the theoretical literature [42, 36, 12, 30], which have shown that effective models can indeed be constructed in some cases, and may be quite different from the dynamics known at small scales. Being phenomenological, such models often cannot easily be interpreted in terms of concrete mechanisms or instantaneous demographic events. They generally do not exhibit the intuitive properties of classical functional responses [41]. But when successful, they can provide useful predictions. The present study should be viewed as exploratory, investigating one hypothetical way of describing large-scale, long-term dynamics by a simple set of effective differential equations.

An important question remains: how can we parameterize our effective model from observational data, collected within or across ecosystems? Indeed, the empirical relations between predation rates and densities of predators and prey do not necessarily translate directly to an effective functional response, i.e. a true dynamical dependence between these three quantities. Other latent factors could be varying between measurements, and creating spurious relations. Nevertheless, we find hints that such confounding variation is weaker than the dependence that we wish to capture. We therefore assume that the observed scaling law can be inserted directly into a simple dynamical model, in order to explore its potential ecological consequences.

The empirical scaling exponents occupy a special position in our model, as they correspond to a maximum in predator abundance and some metrics of stability. While our sublinear model suggests that predation frequency is much lower than in a classic Lotka-Volterra model, it also leads to larger standing predator populations. This apparent paradox is closely related to the “hydra effect”, a term that originally applies to situations where increasing individual predator mortality leads to larger predator populations [2, 51], but can be extended to any situation, such as ours, where consumers that seem individually less fit (e.g. consume less prey than their competitors) may become more abundant in the long term [1].

A second unexpected consequence is the emergence of consistent scaling relationships between predator and prey densities, and between production and biomass within each trophic level. Such scaling laws are widely found in data and postulated in ecosystem models [39, 30]. They are notably used in contexts of ecosystem management, such as fisheries [15, 17]. But these relationships do not emerge spontaneously from arbitrary dynamical models, and only prevail under particular ecological conditions [9]. Our fitted exponents recover simple proportionality rules that are usually associated with donor control.

Our work raises the possibility that consumptive saturation and interference may happen at all scales, not only the classic scale of individual predator behavior [32], and that this prevalence is tied to the empirical robustness of various allometric relationships, and the maximization of predator abundance and persistence.

## 2 Methods

### 2.1 Dataset

We compiled data from 32 observational studies (details in Appendix S4) that reported kill rates *k*_12_ (number of prey killed per predator per year) in the field, as well as prey density *N*_1_ and predator density *N*_2_. For each species, we also collected typical body mass measurements *w* (in kg) and mass-specific metabolic rate measurements *m* (in W/kg) from the AnAge database [20] and two studies by White and Seymour [58] and Makarieva, Gorshkov, Li, Chown, Reich, and Gavrilov [40], as these traits were rarely reported for the populations studied in the field [16]. Whenever multiple values were found, we used the average values for each species.

The final dataset comprises 46 predator-prey pairs including 26 predator species and 32 prey species. Predator species belong to classes Mammalia, Aves and Reptilia, while prey species belong to classes Mammalia, Actinopterygii, Aves and Malacostraca.

For each predator-prey pair, we computed the rate of biomass loss in the prey population due to predation (in kg/km^2^/year) as

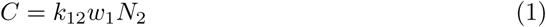

and population biomass densities *B*_1_ = *w*_1_*N*_1_ and *B*_2_ = *w*_2_*N*_2_ in kg/km^2^.

### 2.2 Food chain model

We investigate a simple food chain model of arbitrary length, where we follow the biomass density *B*_*i*_ of trophic level *i* as it changes through time,

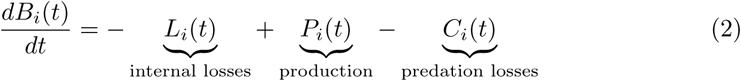

where *L*_*i*_ and *P*_*i*_ represent internal losses and biomass production at level *i*, while *C*_*i*_ represents losses from predation by level *i* + 1. Production at level 1 arises from autotrophic growth

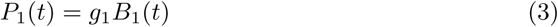

while at higher levels, we assume

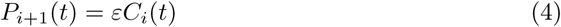

where *ε* (taken here to be constant for simplicity^1^) is the conversion rate between the biomass lost through predation at level *i* and produced at level *i* + 1. Internal losses can arise from individual metabolic costs and mortality, *µ*_*i*_, and from self-regulation or density-dependent processes such as competition and pathogens, *D*_*i*_,

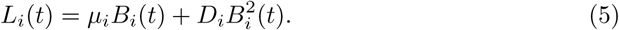

Alternatively, we later consider the possibility of a power-law expression 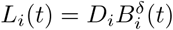 which can interpolate between these two types of losses.

In the following, we only consider stationary properties of the dynamics (2), and thus drop the time-dependence of all quantities, which are assumed to take their equilibrium values.

### 2.3 Functional response

Throughout this study, we consider phenomenological functional responses, emerging at the landscape level from unknown, potentially complex, underlying spatio-temporal dynamics [12]. Consequently, these functions may not follow traditional expectations and intuitions for mechanistic functional responses [41] – for instance, they might not scale additively with the number of prey or predator individuals or species.

While our data is always measured over large spatial scales compared to individual organism sizes, it still covers multiple orders of magnitude in spatial extension and biomass density. This can inform our choice of mathematical expression for the functional response. Indeed, when considering phenomena that range over multiple scales, there are two common possibilities: either there is a scale beyond which the phenomenon vanishes (for instance prey-dependence simply saturates past a critical density), or the phenomenon remains important at all scales. The second possibility drives us to search for a “scale-free” mathematical expression, such as a power-law, which does not truly saturate but exhibits significant variation at all possible scales [10].

Following this argument, we choose here to focus on a “scale-free” power-law dependence of predation losses (see also discussion in Box 1)

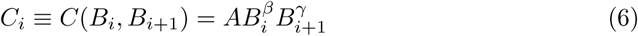

with *β, γ* ∈ [0, 1], and *A* a constant which contains the attack rate, i.e. the basic frequency of encounters for a pair of predator and prey individuals. This attack rate may in principle be specific to each pair of species, while we assume that exponents *β* and *γ* are system-independent.

In this expression, *β* = *γ* = 1 recovers the classic mass action (Lotka-Volterra) model. An exponent *β <* 1 is our counterpart to the well-studied phenomenon of *prey saturation*: predation does not truly saturate here (as the expression is scale-free, see above), but it increases sublinearly with prey density, meaning that the ability of predators to catch or use each prey decreases with their availability [32]. On the other hand, *γ <* 1 indicates *predator interference*: larger predator densities lead to less consumption per capita [53]. In the following, we will consider various arguments suggesting that empirical exponents may instead approximately satisfy the relationship *β* + *γ* ≈ 1, which would imply a simpler ratio-dependent form

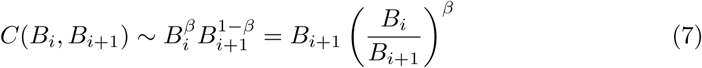

This relationship requires a strong effect of either or both interference and saturation.

These two predation-limiting phenomena are commonly reported and have been studied with various mathematical models. But classic functional responses generally go against our expectation of a “scale-free” expression: they often assume that some density-dependence vanishes when density exceeds a particular scale or threshold. This may be realistic when depicting a given small-scale experimental system, but it is not adequate for data such as ours, compiled from different systems and scales. For instance, we can consider the standard saturating (Michaelis-Menten or Holling Type 2) response [32] and its extension, the DeAngelis-Beddington model [13, 21] with both saturation *H* and interference *I*

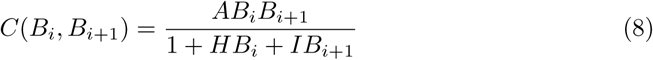

where *H/A* is traditionally defined as the handling time. This expression is applicable to short-term feeding experiments, where consumption is observed for a particular predator-prey species pair whose densities are imposed. The parameters *H* and *I* are contextual: they are expected to depend on species traits [44], but also on the spatial structure and area of each study site. We discuss in Results and Appendix S4 the difficulties associated with using this expression in a cross-ecosystem context.

### 2.4 Links between statistical and dynamical laws

We seek to identify the dynamical relationship *C*(*B*_1_, *B*_2_) between predation losses and the biomasses of predator and prey in empirical data. In our theoretical analysis, we also discuss model predictions for another important observation, the scaling relations between production *P* and biomass *B* within and between trophic levels [30].

Yet, the problem of relating cross-ecosystem statistical laws to underlying dynamical models is subtle, due to the potential covariance between variables and parameters. As a simple example, consider the scaling between production and biomass in one trophic level, assuming that the underlying dynamical law is *P* (*B*) = *rB* with *r* a fixed growth rate. When we consider many ecosystems at equilibrium, each with a different growth rate, their values of *r* and *B* will likely be correlated. If they covary perfectly, the apparent trend will be *P* ∼ *B*^2^ and will not be representative of the true dynamical law.

This issue can be made mathematically explicit in the simple case of an “ecosystem” made of a single species following logistic growth:

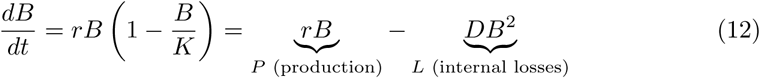

When multiple ecosystems follow the same equation with different parameters, they will reach different equilibria. To avoid confusing the variation within one ecosystem and between different ecosystems, we denote here the equilibrium values in each ecosystem *k* by *B*^(*k*)^, *P* ^(*k*)^ and *L*^(*k*)^ (which satisfy *P* ^(*k*)^ − *L*^(*k*)^ = 0). We can imagine two scenarios that lead to different scalings across systems:

- If systems have the same *r*^(*k*)^ = *r* but different *D*^(*k*)^ (and hence different equilibria *B*^(*k*)^), the relationship *P* (*B*) = *rB* within each system imposes the linear relationship *P* ^(*k*)^ ∼ *B*^(*k*)^ across systems,
- If *r*^(*k*)^ changes across systems while *D*^(*k*)^ = *D* is constant, the relation *L*(*B*) = *DB*^2^ and the equilibrium condition *P* ^(*k*)^ = *L*^(*k*)^ within each system impose the quadratic relationship *P* ^(*k*)^ ∼ (*B*^(*k*)^)^2^ (in other words, we have a perfect colinearity *B*^(*k*)^ ∼ *r*^(*k*)^ across systems)

This example highlights the importance of knowing which parameter (here *r* or *D*) is most system-dependent to predict the empirical cross-ecosystem scaling law, and to identify how it differs from the true within-ecosystem dynamical relationship *P* (*B*) = *rB*.

The same issues occur for all observational scalings, including the scaling of predation losses with prey and predator densities: if we try to fit the relationship 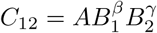 in, we may encounter the problem of correlations between attack rates and population densities. We now discuss how to address this problem in the context of an empirical estimation of *β* and *γ*.

#### Box 1

**Dimensional analysis and functional response**

The difficulty of building a phenomenological cross-scale functional response is that it should account for both:

- between-systems parameter differences (e.g. different study areas, spatial structures, species traits),
- the within-system range of dynamical variables *B*_*i*_ that can lead to saturation and interference if they cross system-dependent thresholds.

We first appeal to dimensional analysis [37]. From (2) we see that predation losses *C*_*i*_ have the dimensions of biomass density over time, e.g. kg/km^2^/year. A classic nondimensionalization choice [62] is to express these losses using predator density and mass-specific metabolic rate (see below for other options)

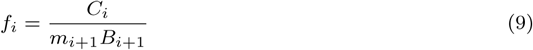

where *f*_*i*_ is the normalized functional response, incorporating all biological mechanisms. Since *f*_*i*_ has no dimension, it can only depend on dimensionless quantities. If we assume that all variation in predation is determined solely by species densities and metabolic rates, we can only construct two dimensionless ratios *π*_1_ and *π*_2_ and *f*_*i*_ must take the form

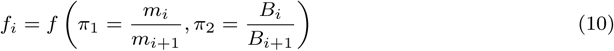

with some arbitrary function *f*. The choice *f* (*π*_1_, *π*_2_) = 1 leads to consumer dependence (*C*_*i*_ = *m*_*i*+1_*B*_*i*+1_), while *f* (*π*_1_, *π*_2_) = *π*_1_*π*_2_ leads to donor dependence (*C*_*i*_ = *m*_*i*_*B*_*i*_). More generally, the expression (10) is strongly reminiscent of ratio-dependent functional response [6], but it does not posit a specific functional form.

To deviate from this ratio-dependent expression, we must use additional parameters to construct other dimensionless quantities. Setting aside metabolic scaling for now, the functional responses (6) and (8) in the main text involve *B*_*i*_ and *B*_*i*+1_ separately, suggesting the form

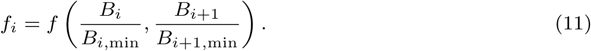

Indeed, it is plausible for nonlinear density-dependence in *f* to involve system-specific thresholds *B*_*i*,min_ (see discussion in Appendix S1). These thresholds can be derived from other parameters that characterize each system, such as movement range or body mass. A further possibility, not investigated here, is that other biological rates may also intervene if metabolic rate cannot be used as a universal “clock” for trophic processes [26].

Dimensionless ratios can capture important differences between systems, and a number of heuristics and mathematical theories exist to guide their choice [10]: for instance, it is often useful to find ratios that are significantly larger than 1 in some systems and smaller than 1 in others, as this can indicate qualitatively different regimes. Thresholds such as *B*_*i*,min_ are commonly found in saturating functional responses [32], where the density-dependence vanishes above a certain scale. If instead we expect a density-dependence that persists over a wide range of scales, we can use a “scale-free” expression for the function *f*, such as a power-law [10].

### 2.5 Empirical analysis

The empirical identification of a relationship between predation losses and species densities such as

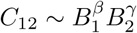

can be affected by two sources of error: the colinearity between the two variables *B*_1_ and *B*_2_, and the colinearity of either of these variables with other factors that may appear in the full expression of *C*_12_ (e.g. the attack rate *A* defined above).

The first problem is that the colinearity between *B*_1_ and *B*_2_ (Fig. 1d) may obscure their respective contributions to *C*_12_ = *C*(*B*_1_, *B*_2_). We can overcome this difficulty by performing a commonality analysis, which is a statistical test based on variance partitioning. We use the function *regr* of the R package *yhat* to check the existence of suppression i.e. the distorsion of regression coefficients due to the colinearity between predictors [49]. This test shows the absence of suppression, allowing us to directly interpret the respective contributions of *B*_1_ and *B*_2_ to *C*(*B*_1_, *B*_2_) below.

**Figure 1.**
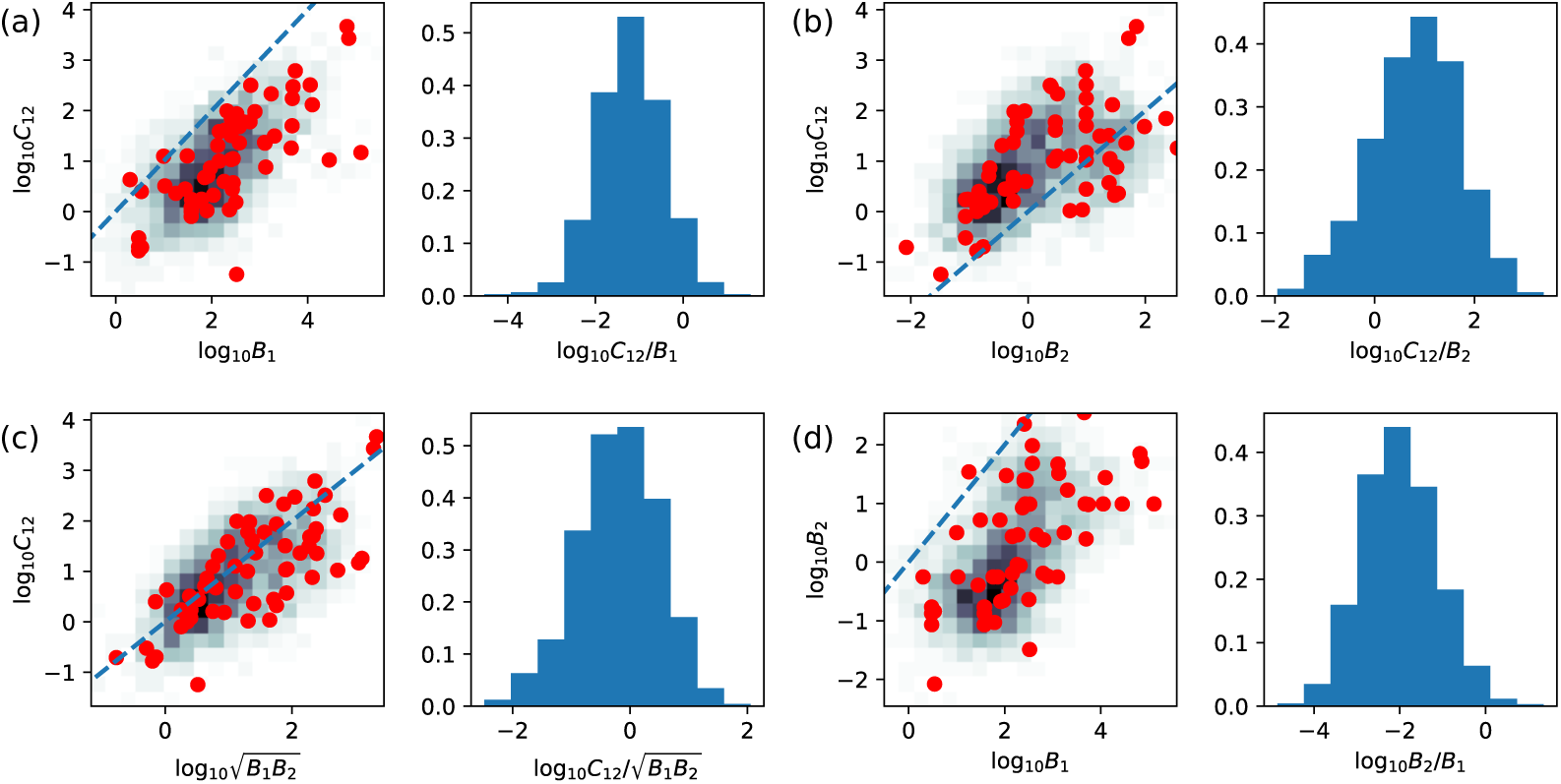
Empirical and simulated scaling relationships betwen biomass densities *B*_1_ and *B*_2_ and predation losses *C*_12_ in logarithmic scale. We show two panels for each pair among (a) *C*_12_ and *B*_1_, (b) *C*_12_ and *B*_2_, (c) *C*_12_ and 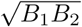, and (d) *B*_2_ and *B*_1_. **Left panels:** Each point corresponds to one study in our meta-analysis, and the dashed line represents the 1:1 relationship. The background color indicates the density of simulated realizations of our dynamical model, retaining only realizations that approximate the empirical distribution (see Sec. 2.5.2). **Right panels:** Histogram of ratio between the two quantities in our simulated ecosystems.

A more important difficulty stems from correlations with other parameters, as suggested in the previous section. If we suppose that, in each ecosystem *k*, predation losses follow

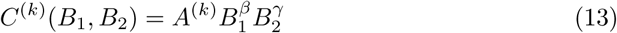

where we are given no information on the species-dependent parameter *A*^(*k*)^, we can only estimate *β* and *γ* from a simple empirical fit if *A*^(*k*)^ is weakly correlated with the observed values of *B*_1_ and *B*_2_. This could be the case if *A*^(*k*)^ varies less between systems, or has less impact, than other factors that control *B*_1_ and *B*_2_, such as primary production or mortality (or factors ignored in our model, such as spatial fluxes, out-of-equilibrium dynamics, and additional interactions).

We employ two distinct methods to identify the exponents *β* and *γ* in our proposed power-law scaling relationship.

The first approach is a direct fit, with the aim of minimizing residual variance: given that *C, B*_1_ and *B*_2_ span multiple orders of magnitude, any combination of these variables that exhibits much less spread suggests an interesting regularity, even though the precise values of the exponents may be incorrectly identified due to colinearities and should be interpreted with care.

The second approach is an attempt to account for at least some unknown factors that may undermine a naive parameter estimation. We focus on the dynamical food chain model (2) and the power-law response (6). We then simulate this model for many random combinations of parameter values (including exponents, but also attack rate, growth rate, mortality, etc.). We only retain model realizations that give triplets (*B*_1_, *B*_2_, *C*) sufficiently close to those found in our empirical data, giving us a distribution of exponents compatible with this evidence, as well as an estimation of how other parameters may affect observed species densities and predation rates.

#### 2.5.1 Direct fit

We note that the factor *A*^(*k*)^ is not unitless. This may be problematic when fitting data collected at different scales. Following the discussion in Box 1, we propose to expand *A*^(*k*)^ in a way that accounts for metabolic scaling and nonlinearity thresholds, i.e.

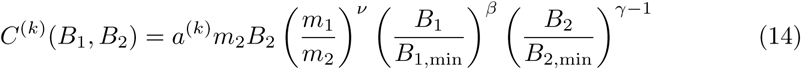

where *m*_2_*B*_2_ sets the dimensions of *C* and all other factors are dimensionless. In particular, *a*^(*k*)^ is an ecosystem-dependent parameter that can be interpreted as a random effect. We will see in the Results section that many terms in this complex equation appear to have a negligible contribution, leading back to our simple theoretical expression (6).

We have no direct measurement of the reference densities *B*_*i*,min_, which are necessary to construct dimensionless ratios, and which we interpret as thresholds for nonlinearity (saturation or interference). Therefore, we must make some assumptions on these values, guided by biological intuitions. The simplest possible assumption is that these thresholds vary independently of other parameters, and can be included into the random effect *a*^(*k*)^. As a second option, if these density thresholds were proportional to individual body masses, *B*_*i*,min_ ∼ *w*_*i*_, the relevant variables for our fit would be population densities *N*_*i*_ rather than biomass densities. A third option, *B*_*i*,min_ ∼ 1*/m*_*i*_, would instead suggest focusing on *E*_*i*_ = *m*_*i*_*B*_*i*_ which has commonly been interpreted as an “energy” density [19]. For each of these choices, the exponents *ν, β* and *γ* may then be identified by a least-squares linear regression in log space. To identify the correct definition of *B*_*i*,min_, we test all three possibilities, and ask which one leads to the best predictions, i.e. the lowest amount of residual variation in *a*^(*k*)^.

#### 2.5.2 Dynamical model estimation

Using our dynamical model (2) with (5) and (6), we generate predator-prey pairs with different parameter values drawn from a broad prior distribution, and compute their equilibrium biomass densities *B*_1_ and *B*_2_ and predation losses *C*_12_. We then retain only model realizations for which the three values (*B*_1_, *B*_2_, *C*_12_) are close to those observed in our data. We repeat this process until we have retained 2500 model realizations, and use the parameter values of these realizations to compute a joint posterior distribution for all our parameters, in particular the exponents *β* and *γ* that we wish to identify.

For two species, our model has 8 parameters, which we draw independently in each model realization. The exponents *β* and *γ* are drawn uniformly in the range [0, 1.5] in order to confirm that they are expected to be less than 1 (the Lotka-Volterra or mass action limit). Other parameters are drawn uniformly on a log scale, over a span of 10 orders of magnitude: the attack rate *A* ∈ [10^−5^, 10^5^], prey productivity *g*_1_ ∈ [10^−4^, 10^6^], predator mortality *µ*_2_ ∈ [10^−8^, 10^2^], and prey and predator self-regulation *D*_1_, *D*_2_ ∈ [10^−5^, 10^5^]. Finally, the biomass conversion efficiency is drawn in the realistic interval *ε* ∈ [10^−2^, 1], see discussion in Barbier and Loreau [9].

We retain only model realizations in which both species survive, and filter the remaining according to their similarity to empirical data. To do so, we train a Kernel Density Estimator (KDE, from the Python library *scikit-learn* using classes *GridSearchCV* and *KernelDensity*) on triplets (log_10_ *B*_1_, log_10_ *B*_2_, log_10_ *C*_12_) in the empirical data. We then compute the same triplet in model realizations, and its score *s* given by the KDE. We retain each model realization with probability 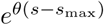, given *s*_max_ the highest score given among empirical triplets. We choose *θ* = 2 as we find empirically that it gives the best agreement between simulations and data for the variance and range of *B*_1_, *B*_2_, *C*_12_ and their ratios.

Finally, we compute the posterior distribution of parameters, i.e. histograms of the parameter values found in accepted model realizations, and study their correlations with each other and with dynamical variables *B*_1_, *B*_2_ and *C*_12_. For additional tests in Fig. 5 we ran the same procedure while restricting which parameters vary (sampling only *g*_1_ and exponents *β* and *γ*, or keeping *D*_*i*_ or *A*_12_ constant) in order to demonstrate the relationship, discussed in Sec. 2.4, between dynamical and statistical scaling laws.

**Figure 2.**
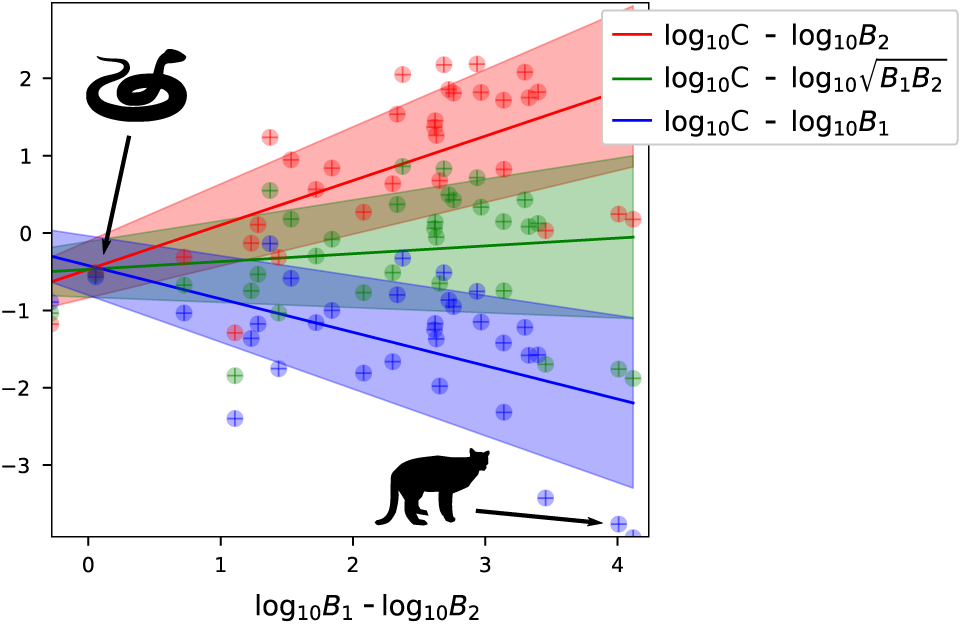
Residual variation with log_10_ *B*_1_*/B*_2_ when assuming strict donor dependence (*C* ∼ *B*_1_), a symmetric square-root law 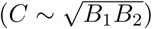, or strict consumer dependence (*C* ∼ *B*_2_). Residues around the square-root law show no trend (*p* = 0.4, *R* = 0.08), and have minimal variance (see Table 1).

**Figure 3.**
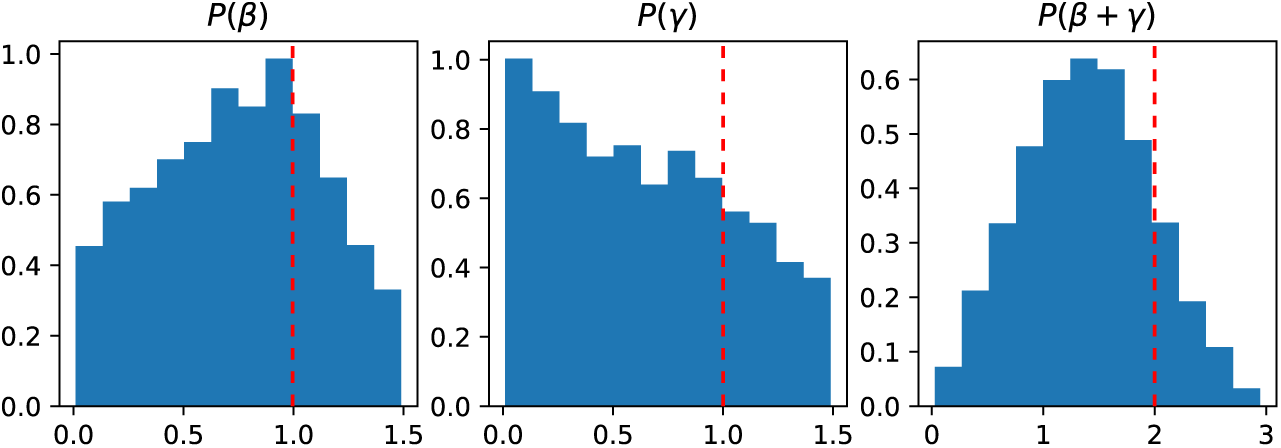
Posterior distributions of exponents *β* and *γ* in power-law functional response (6) for simulations selected to reproduce the empirical distribution of densities *B*_1_ and *B*_2_ and predation losses *C*_12_ (Fig. 1). The prior distribution was uniform over the interval [0, 1.5]. The posterior distributions remain broad, showing that exponents are only weakly constrained by our empirical evidence (with almost no correlation between exponents, *R* = 0.11). The most likely values are *β* ≈ 0.9, *γ* ≈ 0 which approximate the classic model of linear donor-dependence (Box 2), while the median values are more symmetrical, *β* ≈ 0.8, *γ* ≈ 0.5 and hence *β* + *γ* ≈ 1.3. Both show significant departure from the mass action (Lotka-Volterra) hypothesis *β* = *γ* = 1 (dashed lines).

**Figure 4.**
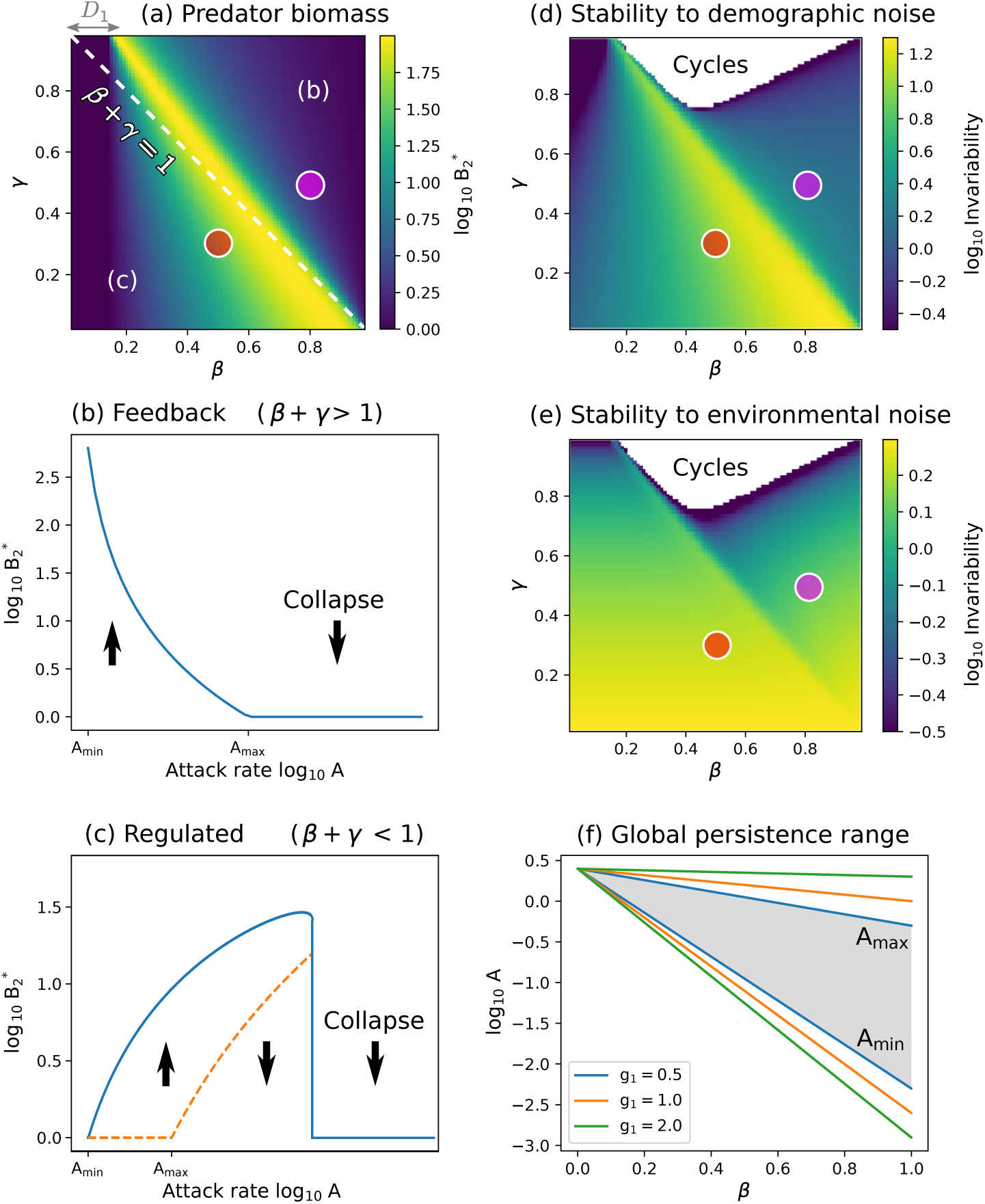
Predator abundance and persistence in our dynamical model (*D*_*i*_ = 10^−3^ at both levels, *g*_1_ = 1, *µ*_2_ = 2, *ε* = 0.8, setting the extinction threshold at *B*_*i*_ = 1). **(a)** Predator biomass density 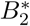 (in log scale) varies with exponents *β* and *γ* in the power-law functional response (6). It is maximized on a boundary separating two different regimes (b and c). This boundary is close to the dashed line *β* + *γ* = 1, but deviates due to self-regulation *D*_*i*_. **(b)** For *β* + *γ >* 1, the resource is depleted by predation (feedback regime in Box 2), and predator population decreases with attack rate, going extinct at *A*_max_. **(c)** For *β* + *γ <* 1, predator biomass (solid line) increases with attack rate, since the resource is mainly limited by its own self-regulation. The predator may survive if *A > A*_min_, but an unstable equilibrium (dashed line) emerges at *A > A*_max_, under which populations can go extinct. **(f)** For the predator to persist in both regimes and for all initial conditions, we must have *A*_min_ *< A < A*_max_ (shaded region). This interval expands with higher *β* (less saturation) and prey growth *g*_1_. **(d**,**e)** Predator invariability (1/coefficient of variation, see Appendix S1) in response to (d) demographic fluctuations and (e) environmental perturbations on the predator. For large *γ* and *β <* 1, the equilibrum can be unstable and replaced by limit cycles. This area is left blank, but invariability remains well-defined in simulations. **(a**,**d**,**e)** The red dot shows the exponents from direct regression *β* ≈ 0.5, *γ* ≈ 0.3, and the purple dot shows median values from our dynamical model fit, *β* ≈ 0.8, *γ* ≈ 0.5.

**Figure 5.**
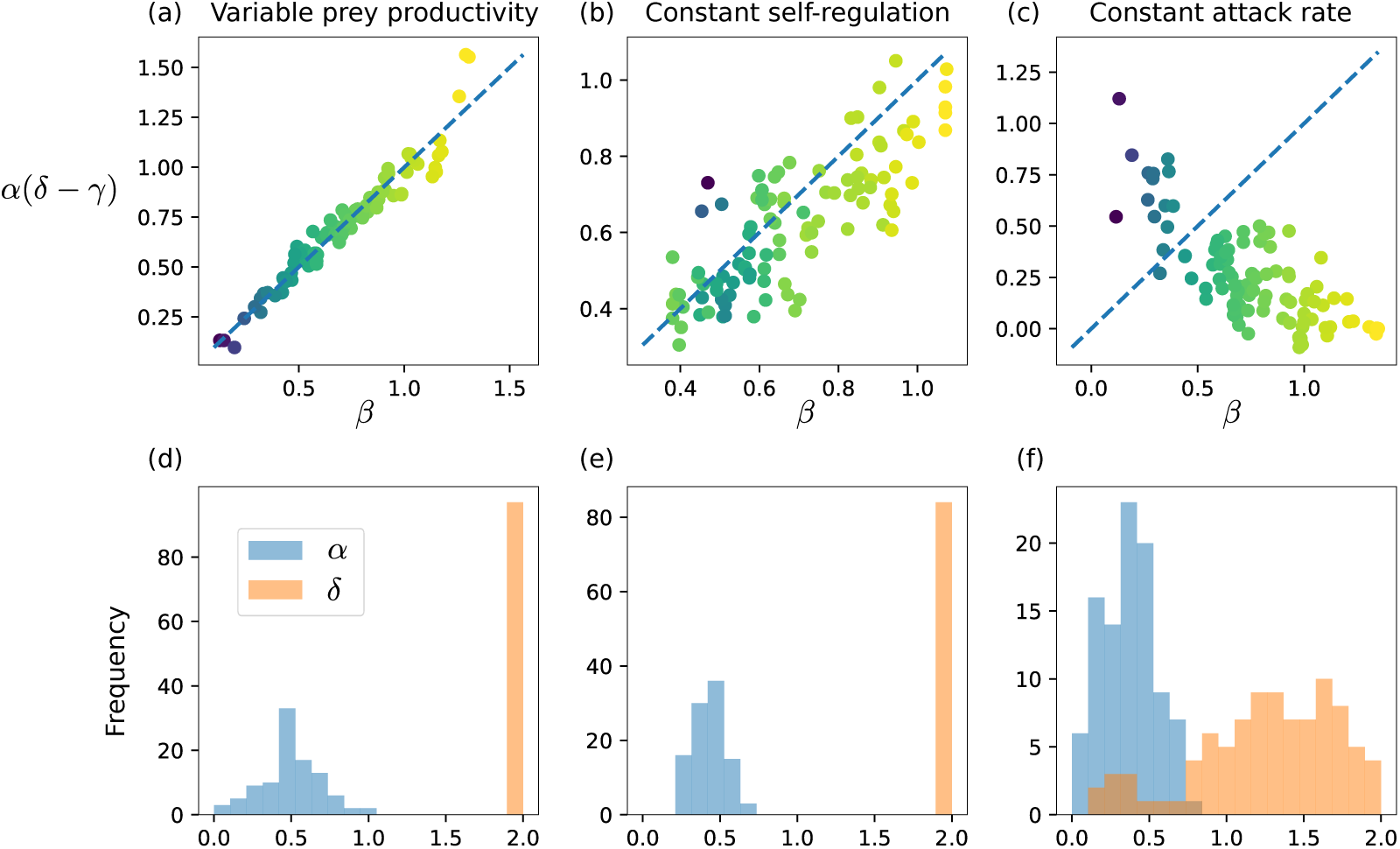
Numerical test of the theoretical relation between scaling laws, *α*(*δ*−*γ*) = *β* in (22), derived in the example of constant self-regulation coefficients *D*_*i*_ across ecosystems. We vary exponents *β* and *γ* in the power-law functional response, and perform regressions to compute the exponents of empirical scaling laws: *α* for the predator-prey density scaling, and *δ* for the prey’s production-density scaling. (a,d) Simulations that differ only through their prey productivity *g*_1_ and exponents *β* and *γ*. (b,e) Simulations where all parameters vary except self-regulation coefficients *D*_1_ and *D*_2_. (c,f) Simulations where all parameters vary except attack rate *A*_12_. We see that the predicted scaling laws depend on which parameter is expected to vary most, so that different calculations may need to be performed depending on which features play the largest role in differentiating observed ecosystems.

## 3 Results

### 3.1 Meta-analysis of kill rates

#### 3.1.1 Results from direct fit

The full expression to be fitted (14) breaks down predation losses *C*_12_ into three different contributions: the observed species densities *B*_*i*_ raised to the exponents *β* and *γ*; the metabolic rates *m*_*i*_, which are often assumed to define the time scale for predation processes; and the unknown typical density scales *B*_*i*,min_, which we interpret as thresholds for nonlinearity (e.g. the scale for interference or saturation).

Our first result is that there does not seem to be a systematic effect due to typical density scales *B*_*i*,min_. As explained in Sec. 2.5.1, our main three choices for these parameters correspond to using biomass densities *B*_*i*_, number densities *N*_*i*_ or “energy” densities *E*_*i*_ = *m*_*i*_*B*_*i*_ as our fitting variables. To showcase these different options, we detail in Table 1 the residual standard deviation for various examples of power-laws involving these variables. These examples are only a few representative points in our systematic exploration of (14), but they illustrate our finding that number densities give the poorest results, while biomass or energy densities give comparable results. For simplicity, we then focus on biomass densities *B*_*i*_, corresponding to the assumption that the reference density *B*_*i*,min_ is independent of body size or metabolism^2^.

**Table 1.** Residual standard deviation for ratios of predation losses and biomass density *B*_*i*_, population density *N*_*i*_, or “energy” density *E*_*i*_ = *m*_*i*_*B*_*i*_. See Fig. 2 for a visualization of the residual variation with *B*_1_*/B*_2_ for the first three expressions. For the rightmost expression, we use the best-fit exponents *β* = 0.47, *γ* = 0.30, and see that this contributes only a small reduction of residual variation compared with the square root expression (second column).

| Expression $y$ | $\frac{C}{B_1}$ | $\frac{C}{\sqrt{B_1 B_2}}$ | $\frac{C}{B_2}$ | $\frac{C}{N_1}$ | $\frac{C}{\sqrt{N_1 N_2}}$ | $\frac{C}{N_2}$ | $\frac{C}{E_1}$ | $\frac{C}{\sqrt{E_1 E_2}}$ | $\frac{C}{E_2}$ | $\frac{C}{B_1^\beta B_2^\gamma}$ |
| --- | --- | --- | --- | --- | --- | --- | --- | --- | --- | --- |
| $std(\log_{10} y)$ | 0.90 | **0.77** | 0.97 | 1.6 | 1.5 | 1.5 | 0.93 | **0.77** | 0.9 | **0.74** |

Our second result is the limited impact of metabolic rates on predation losses. From dimensional analysis (Box 1), we expected a baseline scaling *C*_12_ ∼ *m*_1_ or *m*_2_. But we find that including *m*_*i*_ as a prefactor has no significant impact on any of our parameter estimates. Furthermore, irrespective of our choice of variable, our estimates for exponent *ν* in (14) are all of order *ν* ≈ 0.1, suggesting a weak and perhaps spurious dependence in metabolic rates. We therefore eliminate *m*_*i*_ from our expression (14). We performed a similar analysis (not shown here) using body masses *w*_*i*_, with comparable results.

Given these two simplifications, we are left with the simpler relationship (6) i.e.

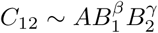

to be fitted. We find the best-fit exponents *β* = 0.47 *±* 0.17 and *γ* = 0.30 *±* 0.19. The limited extent of our dataset implies large uncertainty on the exact exponents, but they are starkly sublinear.

We notice that these exponents lay close to the line *β* + *γ* = 1, i.e. a simpler ratio-dependent model *C*_12_ ∼ *B*_2_(*B*_1_*/B*_2_)^*β*^ which we could have assumed for dimensional reasons (Box 1). If we impose this relationship by setting *γ* = 1 − *β*, we find a similar best-fit value *β* = 0.51 *±* 0.12. Due to the colinearity between *B*_1_ and *B*_2_ (*R* = 0.56, Fig. 1d), we might expect any model of this form to provide a comparable fit, ranging from *β* = 1 (donor dependent, see Box 2) to *β* = 0 (consumer dependent). Nevertheless, our colinearity analysis strongly indicates 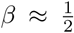 as the most likely value. This is visualized by the fact that *C/B*_1_ and *C/B*_2_ both exhibit a residual variation in *B*_1_*/B*_2_ (*p <* 10^−6^, *R* = 0.61 and *R* = −0.52 respectively, Fig. 2) whereas our best fit exhibits no residual variation (*p* = 0.4, *R* = −0.08), suggesting that neither strict donor nor consumer dependence prevails in our data.

#### 3.1.2 Results from dynamical model estimation

We show in Fig. 3 the results of our second method of parameter estimation, described in Sec. 2.5.2, which consists in simulating our dynamical model with random parameter combinations, and retaining those that produce densities and predation losses comparable to those of the data.

This second approach also supports the idea that *β* + *γ* (whose distribution is centered close to 1) is significantly reduced compared to the mass action expectation *β* + *γ* = 2. However, we notice a stark asymmetry in the posterior distributions of *β* and *γ*: the distribution of *β* has a mode below but close to 1, while the distribution of *γ* is strictly decreasing from 0. This might lend some credence to purely donor-dependent trophic fluxes, a hypothesis which lacked support in our direct fit approach above, see Fig. 2.

We must note that this model exploration is still missing many other factors that may affect *B*_1_ and *B*_2_ and their relationship to *C*_12_, such as non-equilibrium dynamics or interactions with other prey and predator species, among others. These factors may have a greater impact on predators, whose populations are typically smaller, slower to attain equilibrium and more easily perturbed. We could thus expect that measured predator abundances are less strictly related to *C*_12_. This decorrelation may be sufficient to explain the apparent bias toward small *γ*. We discuss below the hypothesis that our results are mainly driven by an artificial donor-dependence imposed by the scale of observation in the data.

#### 3.1.3 Comparison with a classic functional response

The DeAngelis-Beddington (DAB) model (8), or Holling Type 2 with predator interference, can be fitted to our data, but while it may be useful as a mechanistic model for a single system [11], it is unsatisfactory in our macro-ecological approach.

This expression faces the following problem: the thresholds for prey saturation *H* and predator interference *I* are not only particular to each pair of species (e.g. via their body sizes[44]), but they also depend on the spatio-temporal structure and scale of each system. For instance, a day-long experiment in a closed arena may reveal saturation with prey density due to predator handling and satiety. But over a month in an open landscape, a different saturating effect could appear due to prey refuges. If we measure multiple systems at different scales, each may have a different saturation constant, and we cannot fit them all using the same saturating function with fixed, or even independently varying, values of *H* and *I*. We tried to construct some plausible expressions for system-dependent *H* and *I* using other known parameters for each system (e.g. body sizes, metabolic rates and spatial extension), but could not find a successful approach. Assuming *H* and *I* to be constant instead, we perform a fit and find (Appendix S4) that almost all predator-prey pairs are in the saturated part of the function, i.e. *HB*_1_ + *IB*_2_ ≫ 1. This suggests that the DAB model best approximates the data in the limit where it becomes linear in either *B*_1_ or *B*_2_, which is also a special case of our power-law model.

Furthermore, we show in Appendix S4 that the colinearity *B*_1_ ∼ *B*_2_ can make it difficult to differentiate between an additive law, *HB*_1_ + *IB*_2_, and a multiplicative law with exponents adding up to one, 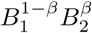, in the denominator of the functional response. This has been put forward as an important criticism of the Cobb-Douglas production function [24] which resembles our power-law model (6). But we find here that prey and predator appear to be both limiting. This requires a fine-tuning of the ratio of handling time to interference, *H/I*, so that *HB*_1_ and *IB*_2_ are comparable in magnitude. This is found in our best fit of the DAB model (8) but is difficult to justify mechanisticially, whereas the same reality may be more clearly and simply encapsulated by our ratio-dependent (*β* + *γ* ≈ 1) expression with *β <* 1.

### 3.2 Theoretical properties and consequences of empirical exponents

#### 3.2.1 Population sizes

We consider two trophic levels, and investigate equilibrium predator density 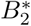 in Fig. 4a as a function of both exponents *β* and *γ*. We observe two distinct regimes: there is only one stable equilibrium throughout the parameter space, but it is dominated either by species self-regulation, or by trophic regulation.

These regimes can be understood by first assuming that self-regulation can be neglected at both levels. In that case, our dynamical model (2) with the power-law functional

##### Box 2

**Definitions**

We introduce two sets of terms to distinguish:

- what sort of dependence is assumed for predation losses *C*, e.g. in our model (6) **donor-dependent predation** A model where *C*(*B*_1_, *B*_2_) = *C*(*B*_1_), e.g. *γ* = 0 **consumer-dependent predation** A model where *C*(*B*_1_, *B*_2_) = *C*(*B*_2_), e.g. *β* = 0 **ratio-dependent predation** A model where *C*(*B*_1_, *B*_2_) = *B*_2_ *C*(*B*_1_*/B*_2_), e.g. *β* + *γ* = 1
- what are the dynamical outcomes of the balance between predation and internal losses

**regulated regime** A solution of our model where each trophic level is mainly limited by self-regulation, as in logistic growth. If we increase the predators’ individual consumption (attack rate *A* and exponents *β* and *γ*), more predators coexist at equilibrium.

**feedback regime** A solution of our model with strong feedbacks between the two trophic levels (each level limits the other). From the predators’ viewpoint, prey stocks are depleted, leading to competition for prey production. If we increase individual consumption, each predator uses up more of that flux, and fewer predators coexist at equilibrium.

These definitions attempt to avoid confusions surrounding the notions of bottom-up and top-down control, which often merge mechanisms (density-dependence) and dynamical outcomes. Intuitively, bottom-up control increases with *β*, top-down control increases with *γ*, and the feedback regime requires *both* types of control to be strong, creating what we previously called antagonistic control [9]. Such mutual limitation is shown in predator-prey cycles, or resource and apparent competition [33].

response (6) has the equilibrium solution (see Appendix S1)

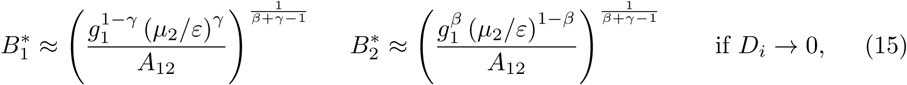

where we see that *β* + *γ* = 1 leads to a singularity, meaning that self-regulation cannot be neglected for these exponents.

The first regime is found when *β* + *γ >* 1, and we call it the *feedback* regime (see Box 2 and Fig. 4b). In this case, the solution (15) is a good approximation at low self-regulation (small *D*_*i*_). We notice that this solution has two counter-intuitive properties: predator biomass *B*_2_ decreases with attack rate *A*_12_ and exponents *β* and *γ*, and increases with predator mortality *µ*_2_ (if there is saturation, *β <* 1). This is because prey stocks are brought to low levels, so predators simply divide between themselves the flux of prey production: the more each predator consumes per capita, the fewer predators can coexist. As we discuss below, this inverse relationship between individual fitness (high attack rate, low mortality) and population abundance has been demonstrated in various models and called the “hydra effect” [2, 51].

If *β* + *γ <* 1, the solution without self-regulation (15) becomes unstable. In the stable equilibrium, the biomass of prey will be mainly determined by their self-regulation i.e. 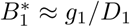, leading to what we call the *regulated* regime (Box 2 and Fig. 4c). In that case, predator biomass

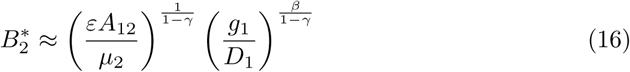

increases with attack rate and with exponents *β* and *γ*. The fact that the previous equilibrium (15) is unstable reveals a type of Allee effect: if initial prey levels are below a threhsold, predators will consume all the prey (consumption will decrease too slowly to adjust to the falling resource level) then collapse.

We show in Fig. 4 that predator abundance is maximized at the boundary between these two regimes. This boundary lies close to the line *β* + *γ* = 1, where predation losses scale as

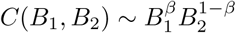

which is close to the exponents suggested by our meta-analysis. We however note that the regime boundary moves away from this line as we decrease attack rates *A*_12_ and increase self-regulation *D*_*i*_ at both levels. For sufficiently high self-regulation, even Lotka-Volterra dynamics can enter the regulated regime, which we interpreted as bottom-up control in previous work [9].

The overarching pattern is that the predator density first increases with attack rate *A*_12_ and exponents *β* and *γ*, when predation losses are still negligible for the prey, up to the regime boundary described above. Past this point, faster consumption leads to significant resource depletion and a drop in predator population [39], as competition increases more than growth.

#### 3.2.2 Dynamical stability

We also observe how functional response exponents affect the stability of predator populations. Fig. 4(d,e) represents the invariability (inverse of the coefficient of variation over time) of predator abundance, computed in two scenarios: demographic stochasticity, where fluctuations (arising from birth and death processes) are proportional to 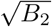, and environmental perturbations which affect the predators proportionally to their abundance *B*_2_, see Arnoldi, Loreau, and Haegeman [7]. Other scenarios and more stability metrics (including asymptotic resilience) are represented in Fig. S1 in Supporting Information, and show qualitatively similar patterns.

We make three main observations. First, the optimal parameter values for stability depend on the perturbation scenario (e.g. demographic or environmental noise) and the choice of stability metric, as having a larger abundance leads to higher stability in some metrics [7]. It is therefore important to know which scenario and metric are relevant for empirical dynamics, and different ecosystems may possibly have different optima. Second, all the cases studied here display a ridge of increased predator stability at the maximum of its abundance, close to the transition line *β* + *γ* = 1, and a drop in stability right after this ridge. Third, while abundance depends almost symmetrically on *β* and *γ*, and is maximized close to the transition between regimes, stability favors systems closer to donor-dependence. Moving toward larger *β* along the line *β* + *γ* = 1 reduces the likelihood of cycles and widens the region of parameters with high stability both to demographic and environmental noise (Fig. 4d and e). More generally, stability to environmental perturbations can be improved by having lower *γ* than what would maximize abundance (i.e. going toward the bottom of Fig. 4e).

### 3.3 Scaling of biomass and production across levels

Previous literature has reported scaling laws between biomass and production within one trophic level,

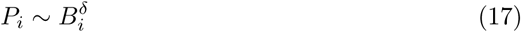

and between biomasses at different trophic levels [30]

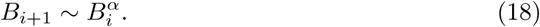

As discussed in Barbier and Loreau [9], neither of these laws can arise for more than two levels in a classic Lotka-Volterra model without self-regulation (*β* = *γ* = 1, *D*_*i*_ = 0).

In the following, we illustrate how such empirical scalings can emerge from underlying dynamical relationships, and how the measured exponents will depend on which parameters drive the cross-ecosystem variation in predator and prey density. There are only two ways in which scaling laws can hold consistently across multiple trophic levels: either they are imposed by a narrow class of functional responses, or they originate from an equilibrium balance between trophic processes and other losses.

The first possibility requires internal losses to be negligible, i.e. *L*_*i*_ = 0 in (2), and predation to be strictly donor-dependent or consumer-dependent (*γ* = 0 or *β* = 0). The functional response then completely determines the scaling between production and biomass,

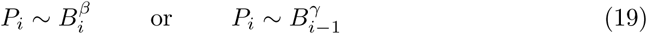

and since *L*_*i*_ = 0,

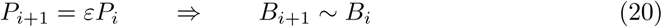

which entails a strict proportionality between predator and prey biomass (exponent *α* = 1). This has been widely discussed by proponents of the ratio-dependent functional response [6], and the existence of such proportionality laws was put forward as evidence of donor control. But since we find nonzero *β* and *γ*, this possibility is excluded by our data.

The second possibility arises in our model with a power-law functional response (6). Following Appendix S3, we illustrate our reasoning in the case where the scaling 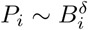 emerges from the density-dependence of internal losses. If we assume that all three terms *C*_*i*_ ∼ *L*_*i*_ ∼ *P*_*i*_ in the dynamics (2) remain of comparable magnitude at equilibrium (i.e. that a finite fraction of biomass is always lost to predation and to internal losses both), the expression of *L*_*i*_ can indeed impose a scaling between production and biomass:

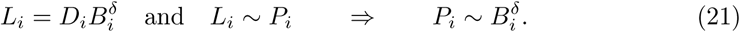

(this requires that the cross-ecosystem variation in densities *B*_*i*_ be mostly due to other parameters, with the exponent *δ* and coefficients *D*_*i*_ varying little between systems, see Sec. 2.4, but a parallel argument can be made when other parameters are held constant). For instance, we have *δ* = 1 for individual mortality, and *δ* = 2 when a self-regulating process, such as intra-specific competition or pathogens, is the main contributor to internal losses. We show in Appendix S3 that a relationship then emerges between the exponents defined in (17) and (18) and the functional response exponents

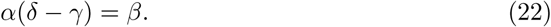

We illustrate this relationship numerically in Fig. 5, using the simulated runs discussed in Sec. 2.5.2 where some parameters are kept constant. We bin simulation runs by their exponents *β* and *γ*, then compute the scaling exponents *α* and *δ* within each bin, by running regressions for the scaling laws (17) and (18). We find that our prediction (22) is successful when self-regulation is constant, and especially when ecosystems differ only by their prey productivity. On the other hand, when self-regulation varies largely, colinearities between parameters and equilibrium densities confound the empirical scaling laws, as discussed in Sec. 2.4, and lead to a violation of this prediction. Similar calculations can however be performed for any assumption about which parameter is mainly responsible for cross-ecosystem differences in predator and prey density.

Commonly assumed scaling exponents for biomass and production are *α* = *δ* = 1, i.e. simple proportionality between predator and prey biomass, and between production and biomass at each level. These relations are used in various mass-balanced models [43] or theory on biomass pyramids [15]. They entail *β* = 1 − *γ*, meaning that the relationship between predation losses and biomasses would be the ratio-dependent expression *C*_12_ ∼ *B*_2_(*B*_1_*/B*_2_)^*β*^ which we have discussed in previous sections.

## 4 Discussion

The predator functional response, describing how predation rates depend on predator and prey abundances, is a crucial component of trophic ecology that has a considerable impact on predator-prey dynamics [32]. Long-standing debates have opposed advocates of various forms of functional response, such as prey-dependent or ratio-dependent expressions [5, 3]. The current consensus seems to be that there is no correct universal expression; rather, different functional responses are appropriate in different settings, and they should be derived mechanistically for each studied ecosystem [11, 3].

From a macro-ecological point of view, however, what matters is how the functional response scales up across several orders of magnitude in abundance, body size and area. Experiments typically measure predation rates for a given pair of species, and how they respond to varying the number of prey and predator individuals, on a short time scale and in an enclosed space. Here, we used kill rates recorded in the field to estimate how predation rates over long times (e.g. a year) vary between species (mainly terrestrial vertebrates) and across much larger spatial scales.

Our empirical analysis points to a phenomenological power-law, which differs from commonly-studied functional responses. If this law does hold dynamically within each ecosystem, it may have striking theoretical implications. The empirical exponents fall close to the boundary between two qualitatively distinct dynamical regimes (approximately the line *β* + *γ* = 1). This boundary has two properties in our food chain model: predator abundance is maximized, and simple allometric scaling laws appear between different ecosystem functions and between trophic levels. We may then ask whether these two important macroscopic properties arise accidentally from a functional response shaped by other constraints, or whether the functional response is in fact selected for its dynamical consequences.

### 4.1 Is the empirical functional response an artefact of donor-dependence?

Our theoretical model suggests that both saturation and interference can lead to similar qualitative consequences for many ecosystem properties. Yet, two main empirical results suggest that reality is closer to classic donor-dependence, i.e. assuming that trophic fluxes are only determined by the amount of resource, and in turn determine the abundance of consumers, and not the reverse. The first clue is the fact that *C*_12_*/B*_1_ is largely below 1, and the second is that exponent *γ* is consistently lower than *β*. As discussed previously, the precise values of these exponents are weakly constrained empirically and could be confounded by unknown latent factors, disguising an underlying process where *β* = 1 and *γ* = 0.

To properly assess the value of these clues, it is important to recognize that donor-dependence is not a necessity. Predation losses are in principle limited not by standing biomass, but by production,

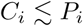

which can be much larger than standing biomass. For instance, a Lotka-Volterra model with a classic biomass pyramid *B*_*i*+1_ ∼ *B*_*i*_ would predict the predation and production of all consumers to scale *quadratically* with biomass, as 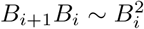. In other words, more and more productive systems would exhibit more and more predation, but populations would only increase as the square root of these fluxes, and prey density *B*_1_ would not represent a bound on predation. A supralinear scaling of production and losses with biomass is not implausible. Analogous relationships have been proposed in economic systems, with empirical exponents ranging from 1.16 (economic growth as a function of city size [14]) to 1.6 (stock trading rate as a function of number of shares [8]). We could also invoke empirical facts that have been interpreted as evidence for the Allee effect [35], since this effect assumes that per-capita growth rate increases with density, and hence, that production *P*_*i*_ is supralinear with biomass *B*_*i*_.

There is, however, an important reason for expecting donor-dependence to prevail, especially at large spatial and temporal scales where many ecological processes are aggregated: the possibility of a *bottleneck* in the dynamics. In various ecological contexts, only a fraction of possible prey (or predators) are available for predation events. This can include the existence of prey refuges in space [47], or the specialization of predators on particular age classes. The rate at which prey become available can be the limiting factor in the overall predation intensity, which therefore becomes purely donor-dependent. For example, if predators target prey of age one and above, these prey must have appeared in the previous year’s census, imposing an insuperable bound *C*_*i*_ *< B*_*i*_ (with a year’s delay, ignored under equilibrium assumptions).

The evidence for or against this possibility is unclear here. In most of the data that we study, *C*_12_ is sufficiently small compared with *B*_1_ that it could still plausibly increase with predator abundance, and there is no systematic preference for mature prey. On the other hand, our estimate of *B*_1_ might miss other factors (such as refuges) limiting prey availability, as we discuss next. While a broader analysis across taxonomic groups, timescales and life histories would be needed to test for possible artefacts such as those suggested above, our provisional conclusion is that this empirical scaling of predation with densities is indeed meaningful, and goes beyond restating the classic donor-dependence hypothesis.

### 4.2 Is this functional response a consequence of biological constraints?

Nonlinear functional responses, with saturation and interference, are generally assumed to be imposed by a broad range of microscopic or mechanistic factors (physiology, behavior, spatial structure, individual trade-offs between competition and consumption, etc). In various ways, these factors limit the space or time available for encounters, leading to less predation than under mass action.

The most immediate candidate mechanism to explain a pattern that holds across ecosystems and scales is spatial structure. For instance, local prey refuges have been proposed as a classic explanation for a donor-dependent [47] or DeAngelis-Beddington response [25]. More generally, spatial structure has been shown to lead to power-law-like functional responses in agent-based or spatially explicit simulations [34, 11]. But this seems to allow a wide range of possible exponents, and therefore, it does not provide an unequivocal explanation for our observations.

A functional response 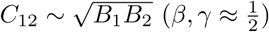, similar to our measurements, could be imposed by spatial structure in two dimensions, if prey and predators occupy almost mutually exclusive regions and only meet at the boundaries (given that the periphery of a region scales like the square root of its area). This could happen due to local prey depletion by predators, a plausible explanation although it requires very strong negative correlations in space [52], and would predict a different scaling in three-dimensional settings [44] which remains to be tested.

We cannot conclude, based on our data alone, that spatial heterogeneity is the main driver of our empirical functional response. Spatially explicit data may be necessary to test this hypothesis. If spatial structure does not impose an exponent, but allows a range of possibilities, it could provide the means through which predators and prey organize to achieve a functional response that is selected by other processes [46, 27].

### 4.3 Can this functional response emerge from selection?

The functional response and exponents found here are close to those which theoretically maximizes predator abundance and some metrics of stability. Yet this comes at an apparent cost to the individual predator, which consumes less here than under a classic Lotka-Volterra (mass action) scenario. We recall the so-called hydra effect [2, 51], a counter-intuitive but common phenomenon in resource-based dynamics: decreasing the fitness of individual consumers, e.g. increasing their mortality or reducing attack rates, can lead to larger populations in the long run, when this reduces competition more than it reduces growth. We showed that a similar effect applies here to consumptive saturation and interference, as represented by the power-law exponents *β* and *γ* in the functional response. Maximal predator abundance is reached for exponents close to the relationship *β* + *γ* = 1, which holds approximately in data^3^.

This resonates with long-standing debates in ecology and evolution, in particular the group selection controversy [59] and its accounts of selection for population abundance [60] or persistence [61]. Should we conclude here that consumption rates are selected to optimize population-level properties? And would this optimization require population-level selection, or can it arise strictly at the individual level? We now show that these are valid possibilities in our setting.

A first possibility is that our proposed functional response is outcompeted at the individual level, but selected at the population level, due to some positive consequences of having larger abundances. Extending our model to include simple adaptive dynamics of attack rate and scaling exponents, we indeed find that this functional response is dominated by other strategies (Appendix S2). We observe maladaptive evolution: mutants with ever faster consumption will outcompete and replace residents, reaching ever lower equilibrium abundances, up until the point where the predators go extinct. A classic solution is to invoke competition between groups in a spatial setting [60, 59], as a larger group can send out more propagules and will often disperse faster [28]. These intuitions pervade a large literature on the evolution of “prudent” predation strategies through group selection, also known as the milker-killer dilemma [56, 45, 46, 27].

A less commonly considered possibility is that our functional response is favored even at the individual level, as a result of direct competitive interactions. The dominance of faster consumers relies on the assumption that competition takes place only through resource consumption. In the presence of other competitive interactions that are not tied to resource levels, a larger standing population can resist invasion both by mutants and by other species. We suggest in Appendix S2 that, for the resident population to be evolutionarily stable, this non-consumptive competition must induce higher mortality in individuals that *search* for more resources, regardless of their success. This feature seems plausible for territorial behavior and aggression as widely displayed by higher vertebrates ^4^, but could also be induced e.g. by allelopathy in other organisms. Under these conditions, slow consumers can avoid being overtaken by fast consumers, even within a single population.

As a further perspective, all our arguments so far have taken the viewpoint of the predator’s strategy and abundance, despite the fact that our empirical results suggests a roughly equal role of predator and prey density in limiting predation. Our best-fit exponents situate real systems close to the limit between a regime where prey suffer from predation, and a regime where they are mainly self-regulated. It is thus possible that prey strategies, or a balance between prey and predator selective pressures, are at play. We conclude that selection on the functional reponse exponents to maximize population size can easily arise in variations of our model, provided that we include non-consumptive processes (from dispersal to aggression). Some of these explanations involve competition between populations, while others take place within a single population, but all favor strategies that maximize predator abundance.

### 4.4 Can this functional response explain other power-law relations between ecosystem functions?

Relations between production and biomass, and between predator and prey densities are widely assumed and observed to take linear or power-law forms [15, 30]. Yet, we have shown here and in previous work [9] that such relations do not emerge universally from arbitrary dynamical models. They require particular ecological conditions, where internal losses *L*_*i*_ within trophic level *i* are comparable to, or larger than, predation losses *C*_*i*_ (both in their magnitude and in their density-dependence). For instance, in a Lotka-Volterra model, strong density-dependent self-regulation within a trophic level is required to observe linear (isometric) relations between different functions and between levels, creating classic biomass and energy pyramids [9]. Alternatively, exponents *β* + *γ* = 1 in our proposed functional response recover the same linear relations and pyramids. When these conditions are not satisfied and predation losses are larger, the predator-prey dynamics can enter a feedback regime (see Box 2) in which prey stocks are exhausted and may not follow any scaling relationship to either prey productivity or predator stocks, a property which has already been proposed as evidence for top-down control of prey by predators [50].

A recent meta-analysis by Hatton, McCann, Fryxell, Davies, Smerlak, Sinclair, and Loreau [30] found sublinear scalings of production with biomass (with exponent *α*) and of predator density with prey density (with exponent *δ*), following 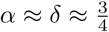, reminiscent of metabolic allometry. Assuming that this cross-ecosystem law also holds dynamically within one system, so that our model results apply, this requires strong interference and saturation, *β* + *γ <* 1. The empirical evidence is too weak to decide whether *β* + *γ* ≈ 1 (leading to isometry) or *β* + *γ <* 1 (leading to allometry), but both possibilities fall within the range of our estimates.

In summary, the existence of well-defined scaling laws across trophic levels, between ecosystem functions such as biomass and production, is less self-evident than it may appear. We suggest here that some empirically-supported laws might be secondary consequences of the mechanisms that determine our phenomenological functional response.

### 4.5 Conclusions

We have shown that the intensity of predation across a range of spatial scales and species can be modeled as a power-law function of both consumer and resource densities, that deviates strongly from the mass action (Lotka-Volterra) assumption. Classic functional responses cannot be automatically assumed to be adequate for macroscopic data: descriptions that are valid at small scales may not always be used at larger scales, as dynamics are driven by very different mechanisms, from individual behavior all the way to landscape heterogeneity. Similar phenomena, such as saturation and predator interference, may nevertheless emerge from these different causes. Our empirical investigation should be extended to other taxonomic groups, and confronted to experimental evidence, to pave the way for a deeper understanding of the emergent functional response at macro-ecological scales.

This phenomenological power-law model can recover classic donor control as a special case, but our empirical estimates are also compatible with a more symmetrical expression where predator and prey are roughly equally limiting, leading to an unusual square root expression. This model is reminiscent of the Cobb-Douglas production function in economics [18], which similarly arises in a macroscopic context, and is also disputed as either a salient empirical fact or a simple consequence of colinearity and aggregation [24]. Nevertheless, the main observation in both fields is that two factors – labor and capital, or consumers and resources – are co-limiting, so that doubling the output requires doubling each of the two inputs.

Scaling laws measured across ecosystems do not always hold dynamically within a single ecosystem [29]: other latent variables could differ between ecosystems, altering the relationship between predation and abundances that we would observe locally. Our results are also limited by the fact that we consider only one interaction at a time, as we generally lack data for other prey and predators interacting with our focal species. Future work should employ data on temporal change to test whether our proposed functional response truly applies to the dynamics of an ecosystem. If it does, we have shown that our empirical exponents have important implications for various features of trophic systems: in particular, they are close to maximizing predator population size and some measures of stability. This could reflect an universal selection pressure acting upon consumption rates. We notably discussed the possibility that non-consumptive competition (e.g. territorial exclusion or allelopathy) would prevent the usual dominance of smaller populations of faster consumers, and offer an explanation for the evolution of strategies that limit predation to maximize predator abundance.

We hope that future work can address two fundamental questions: (1) whether the functional response that we observe stems purely from aggregation or from other ecological phenomena, and (2) whether some universal principle, such as selection for larger populations, could explain this functional response and, through it, the emergence of other widespread macro-ecological scaling laws.

## Supporting information

Simulation code

## Acknowledgments

We thank F. Barraquand, I. Hatton and J. Prunier, as well as the PCI reviewers and recommender, for many helpful comments. This work was supported by the TULIP Laboratory of Excellence (ANR-10-LABX-41) and by the BIOSTASES Advanced Grant, funded by the European Research Council under the European Union’s Horizon 2020 research and innovation programme (666971). Version 4 of this preprint has been peer-reviewed and recommended by Peer Community In Ecology (DOI: 10.24072/pci.ecology.100051).

## Author contributions

All authors contributed to the design of the study and the writing of the manuscript. MB analyzed the theoretical model. LW designed the meta-analysis and assembled the data. LW and MB jointly analyzed the data.

## Data and code accessibility

The dataset and simulation code are available in the online Supplementary Materials, with metadata and additional information in the Supplementary Appendices.

## Footnotes

1 The conversion rate *ε* is known to vary, but its range is limited compared to other quantities which can span orders of magnitude, see discussion in Barbier and Loreau [9]. We choose to ignore ecological contexts in which predator production may be density-dependent even for a fixed amount of consumption, e.g. a reduced offspring number in denser groups, which could be represented by a function *ε*(*B*_*i*+1_).

2 This is consistent with ssuming that the thresholds *B*_1,min_ and *B*_2,min_ are equal or proportional to the minimal sustainable population densities for prey and predators. Cross-species data from Hatton, McCann, Fryxell, Davies, Smerlak, Sinclair, and Loreau [30] and Stephens, Vieira, Willis, and Carbone [54] suggest that the minimal numeric density is roughly inversely proportional to body mass, *N*_*i*,min_ ∼ 1*/w*_*i*_. Therefore, *B*_*i*,min_ = *w*_*i*_*N*_*i*,min_ is expected to be size-independent, and exhibits no known link to metabolism.

3 More precisely, predator abundance is maximized at the boundary between two dynamical regimes, which is given by *β* + *γ* = 1 when species self-regulation is low, but moves toward higher exponents as we increase self-regulation and can be seen in Lotka-Volterra models [9].

4 As an illustration, lions are both outliers in our dataset, since they display lower consumption rates than the general trend (see Appendix S5), and are a major cause of mortality among other carnivores [31]. These two features together may indicate a counter-intuitive predator strategy, which involves both consuming less prey and spending more time in non-consumptive competition, leading to and benefitting from population maximization.

