## Supplementary Appendices for "A macro-ecological approach to predators’ functional response"

### Supplementary Materials: A macro-ecological approach to trophic functional response and its consequences, from population sizes to allometric laws

Matthieu Barbier, Laurie Wojcik and Michel Loreau

March 4, 2020

### S1 Power-law functional response

#### S1.1 Basic model

We study the Lotka-Volterra predator-prey dynamics with a power-law functional response

$$\frac{dB_1}{dt} = \phi_1(B_1, B_2) = g_1 B_1 - D_1 B_1^2 - F(B_1, B_2)$$
$$\frac{dB_2}{dt} = \phi_2(B_1, B_2) = -\mu_2 B_2 - D_2 B_2^2 + \varepsilon F(B_1, B_2) \quad \text{(S1)}$$

where the predation term is given by

$$F(B_1, B_2) = A_{12} \left(\frac{B_1}{B_{1,\min}}\right)^{\beta} \left(\frac{B_2}{B_{2,\min}}\right)^{\gamma} \quad \text{(S2)}$$

and $B_{1,\min}$ and $B_{2,\min}$ are thresholds discussed below, and we assume that the exponents lay in the interval $\beta, \gamma \in [0, 1]$. We refer to Box 1 in the main text for justifications from dimensional analysis.

##### S1.1.1 Exponents and thresholds

Sublinear exponents represent mechanisms limiting the efficiency of predation: saturation with the number of prey (if $\beta < 1$) and consumer interference (if $\gamma < 1$). This functional response only holds if $B_1 > B_{1,\min}$ and $B_2 > B_{2,\min}$, since otherwise we obtain higher predation rate than the Lotka-Voltera functional response, going against the intended meaning of these exponents. These thresholds are therefore analogous to e.g. half-saturation densities in the sense that they indicate when limiting mechanisms become important (we could assume that the functional response becomes Lotka-Volterra below the thresholds, but we never study this limit here).

As a simplest case, we can take $B_{i,\min} = w_i$ the body mass of an individual of species $i$, in which case the ratio becomes $N_i$ the numeric abundance, and the situation $N_i < 1$ entails that the species is extinct. In our theoretical analysis, we follow the option of defining $B_{i,\min} = 1$ as the extinction threshold, assuming that biomasses are measured in the corresponding units.

##### S1.1.2 Metabolic scaling

If we wish to follow the common assumption that predation losses should scale with predator biomass and metabolism [1] with some proportionality constant $a$, we can rewrite

$$A_{12}\left(\frac{B_1}{B_{1,\min}}\right)^{\beta}\left(\frac{B_2}{B_{2,\min}}\right)^{\gamma} = am_2B_2\left(\frac{B_1}{B_{1,\min}}\right)^{\beta}\left(\frac{B_2}{B_{2,\min}}\right)^{\gamma-1} \tag{S3}$$

to see that this is equivalent to defining $A_{12}$ as

$$A_{12} \equiv \frac{am_2}{B_{2,\min}}. \tag{S4}$$

##### S1.1.3 Self-regulation

In the equations above, we distinguish between direct self-regulation, represented by $D_1$ and $D_2$, and predator interference which can cause $\gamma < 1$. Interference is conceptually a type of self-regulation for the predator, but not a direct one (it is conditional on interactions with prey), whereas $D_2 \neq 0$ means that mortality increases (or growth decreases) directly as a result of the existence of conspecifics.

At equilibrium (when an equilibrium exists), there has to be a balance between positive and negative terms in (S1). There is always one positive term and two negative terms, so one of the two negative terms can dominate the other and determine the equilibrium. Thus, self-regulation becomes important for the prey when

$$D_1B_1^2 \gtrsim A_{12}B_1^{\beta}B_2^{\gamma} \quad \Rightarrow \quad B_1 \gtrsim \left(\frac{A_{12}}{D_1}B_2^{\gamma}\right)^{\frac{1}{2-\beta}} \tag{S5}$$

and for the predator, when

$$D_2B_2^2 \gtrsim \mu_2B_2 \quad \Rightarrow \quad B_2 \gtrsim \frac{\mu_2}{D_2} \tag{S6}$$

This just gives orders of magnitude: we know that we can neglect self-regulation as long as $B_1$ and $B_2$ are much smaller than these values.

##### S1.1.4 Stability analysis

The Jacobian matrix at equilibrium $B_i = B_i^*$ is given by

$$J = \begin{pmatrix} \frac{\partial \phi_1}{\partial B_1}(B_1^*, B_2^*) & \frac{\partial \phi_1}{\partial B_2}(B_1^*, B_2^*) \\ \frac{\partial \phi_2}{\partial B_1}(B_1^*, B_2^*) & \frac{\partial \phi_2}{\partial B_2}(B_1^*, B_2^*) \end{pmatrix} \tag{S7}$$

After some calculations [1], we find:

$$\frac{1}{B_1^*}J_{11} = -D_1 - AB_2^*\frac{\partial F}{\partial B_1} \tag{S12}$$

$$\frac{1}{B_1^*}J_{12} = -A\left(F + B_2^*\frac{\partial F}{\partial B_2}\right) \tag{S13}$$

$$\frac{1}{B_2^*}J_{21} = \varepsilon A\left(F + B_1^*\frac{\partial F}{\partial B_1}\right) \tag{S14}$$

$$\frac{1}{B_2^*}J_{22} = -D_2 + \varepsilon AB_1^*\frac{\partial F}{\partial B_2} \tag{S15}$$

The most important feature of this set of equations is the fact that $J_{11}$ can become positive if $\partial F/\partial B_1$ is sufficiently negative ($F$ decreases quickly with $B_1$ i.e. high saturation). This is the main source of instability and cycles, as we can see from a simple stability analysis.

We can compute the eigenvalues as

$$\lambda_{\pm} = \frac{T \pm \sqrt{T^2 - 4D}}{2} \tag{S16}$$

with

$$T = J_{11} + J_{22} \tag{S17}$$

the trace of the matrix and

$$D = J_{11}J_{22} - J_{12}J_{21} \tag{S18}$$

its determinant. Hence the sign of

$$T^2 - 4D = (J_{11} - J_{22})^2 + 4J_{12}J_{21} \tag{S19}$$

[1] First, notice that

$$\frac{dB_i}{dt} = \phi_i(\vec{B}) = B_i\Phi_i(\vec{B}) \tag{S8}$$

with $\Phi_i$ the per-capita growth function, so the Jacobian matrix at equilibrium $B_i = B_i^*$ has off-diagonal terms

$$J_{ij} = \frac{\partial \phi_i}{\partial B_j} = B_i^*\frac{\partial \Phi_i}{\partial B_j} \qquad (i \neq j) \tag{S9}$$

and on the diagonal

$$J_{ii} = \Phi_i(\vec{B}^*) + B_i^*\frac{\partial \Phi_i}{\partial B_j} \tag{S10}$$

but $\Phi_i(\vec{B}^*) = 0$ by definition of the equilibrium, hence the Jacobian is always the equilibrium abundance multiplied by the partial derivative of the per-capita growth function

$$J_{ij} = B_i^*\frac{\partial \Phi_i}{\partial B_j}(\vec{B}^*) \qquad \forall i, j \tag{S11}$$

As a consequence, constant growth/mortality terms such as $g_i$ or $\mu_i$ disappear when taking the derivative and do not intervene explicitly in the Jacobian (they appear indirectly through equilibrium abundances).

determines whether eigenvalues will have an imaginary component, indicating oscillating tendencies. These properties are shown in Fig. S1 (a-d). Stability can also be computed as invariability, see e.g.[2] for technical details. We show predator invariability, computed analytically as $1/CV_2$ (inverse of the coefficient of variation of the predator population), in response to a demographic or environmental perturbation applying either only to the predator or both levels in Fig. S1 (e-h). A positive dominant eigenvalue indicates cycles, in which case invariability remains well-behaved numerically but cannot be computed analytically.

#### S1.2 Low self-regulation equilibrium

##### S1.2.1 Abundances

If we assume that the equilibrium abundance is low enough to neglect self-regulation, then we have

$$0 \approx g_1 B_1 - A_{12} B_1^{\beta} B_2^{\gamma} \tag{S20}$$

$$0 \approx -\mu_2 B_2 + \varepsilon A_{12} B_1^{\beta} B_2^{\gamma} \tag{S21}$$

This admits a solution

$$B_1^* = \left( \frac{g_1^{1-\gamma} \left(\mu_2/\varepsilon\right)^{\gamma}}{A_{12}} \right)^{\frac{1}{\beta+\gamma-1}} \tag{S22}$$

$$B_2^* = \left( \frac{g_1^{\beta} \left(\mu_2/\varepsilon\right)^{1-\beta}}{A_{12}} \right)^{\frac{1}{\beta+\gamma-1}} \tag{S23}$$

We recall that our model is only meaningful when $B_1^*, B_2^* > 1$ (which we take here as our extinction threshold).

We will focus in particular on the condition $B_2^* > 1$, meaning that the consumer can survive. We see that there are two cases: if $\beta + \gamma > 1$, the contents of the parentheses should be larger than 1 for this condition to hold, whereas if $\beta + \gamma < 1$, the contents should be smaller than 1.

For reasons that will become clear later (when comparing this case to equilibria with self-regulation), we choose to define a rescaled attack rate

$$\tilde{A} = \frac{A_{12}\varepsilon}{\mu_2} \tag{S24}$$

and observe that survival of the predator at equilibrium, $B_2^* > 1$, requires

$$\tilde{A} < \tilde{A}_{\max} \equiv \left( \frac{g_1 \varepsilon}{\mu_2} \right)^{\beta} \qquad \text{if} \qquad \beta + \gamma > 1 \tag{S25}$$

Hence, predators with larger attack rates will drive themselves to extinction in this equilibrium.

We will now see that this equilibrium becomes unstable if $\beta + \gamma < 1$, and explain that predators with large attack rates $\tilde{A} > \tilde{A}_{\max}$ therefore exhibit an Allee effect when exponents are low.

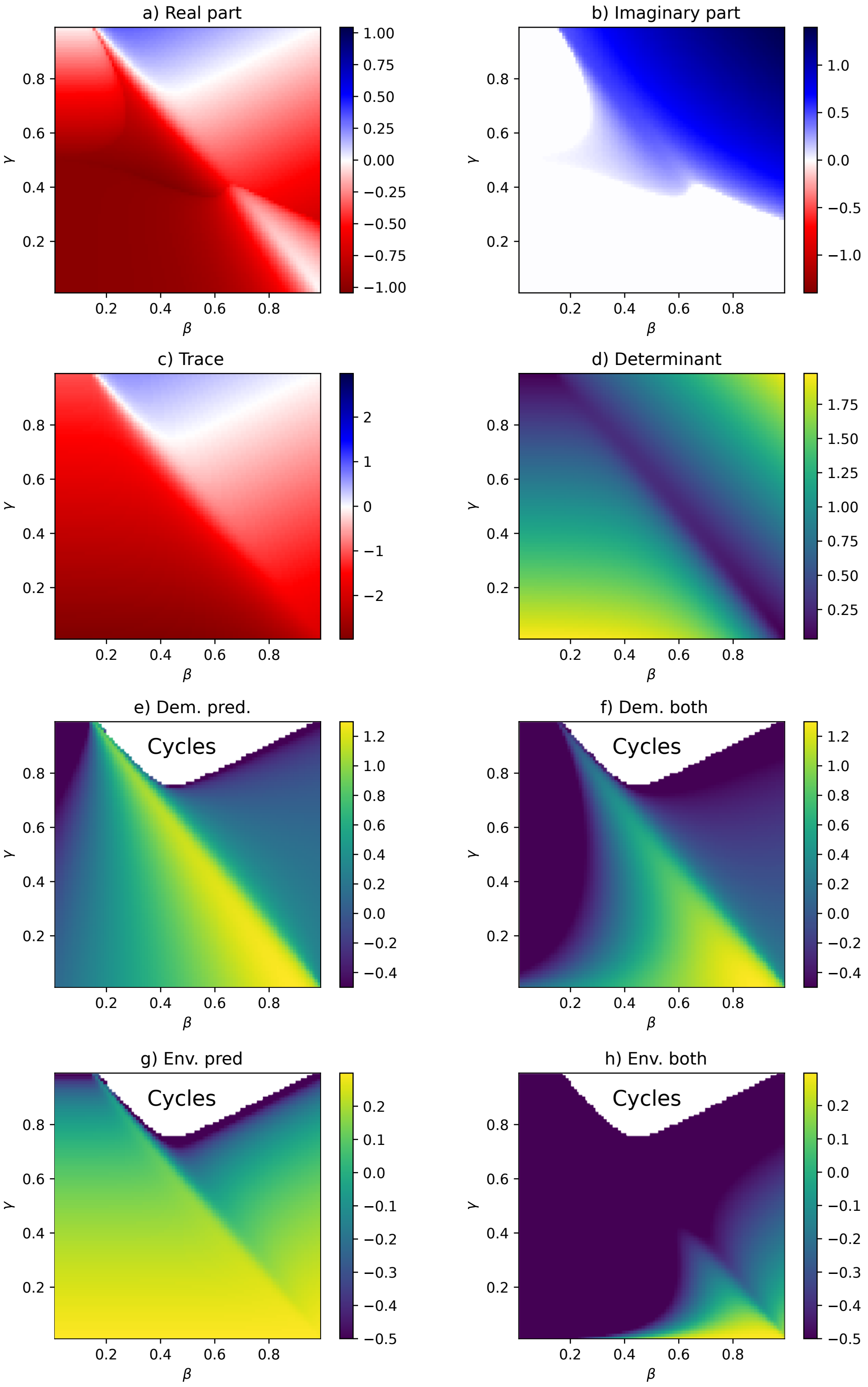

Figure S1: Stability properties of the model, using eigenvalues (a-d) and analytically calculated variability (e-h). (a) Real part of the dominant eigenvalue. (b) Imaginary part of the dominant eigenvalue. (c) Trace $T$. (d) Determinant $D$. (e-h) Predator invariability in response to demographic (e,f) or environmental (g,h) noise applying to the predator only (e,g) or to both levels (f,h).

##### S1.2.2 Stability analysis

If $D_1 = D_2 = 0$, the Jacobian derived in Sec. S1.1.4 takes a simple expression

$$J = \begin{pmatrix} (1-\beta)g_1 & -\gamma\mu_2/\varepsilon \\ \beta g_1 \varepsilon & -(1-\gamma)\mu_2 \end{pmatrix} \tag{S26}$$

We can compute the eigenvalues as

$$\lambda_\pm = \frac{T \pm \sqrt{T^2 - 4D}}{2} \tag{S27}$$

with $T$ the trace of the matrix and $D$ its determinant

$$T = (1-\beta)g_1 - (1-\gamma)\mu_2 \tag{S28}$$

$$D = (\beta + \gamma - 1)g_1\mu_2 \tag{S29}$$

Since $D$ is the product of eigenvalues, we know that if $D < 0$ ($\beta + \gamma < 1$) the system cannot be stable (one eigenvalue is positive and the other is negative).

The system is only stable if $D > 0$ and $T < 0$ (both eigenvalues are negative), i.e.

- $\beta + \gamma > 1$ and
- $\mu_2 > g_1(1-\beta)/(1-\gamma)$

Thus we can have a stable system if predator mortality $\mu_2$ is strong enough.

If $\beta+\gamma < 1$, the equilibrium is unstable, but $D < 0$ and therefore eigenvalues are real: we do not get cycles, but an Allee effect (if abundances start below some threshold, they crash, else they increase until they reach values where we cannot neglect self-regulation – the dynamics without self-regulation simply diverges).

As an aside, we note that oscillations require complex eigenvalues, i.e. $4D > T^2$ in (S27), which we can rewrite using

$$D = (\beta + \gamma - 1)g_1\mu_2 = -(1-\beta)g_1(1-\gamma)\mu_2 + \beta\gamma g_1\mu_2 \tag{S30}$$

$$T^2 - 4D = \left((1-\beta)g_1 + (1-\gamma)\mu_2\right)^2 - \beta\gamma g_1\mu_2 \tag{S31}$$

where we see that oscillations disappear when $\beta$ and $\gamma$ are small enough. In the marginally stable case $\gamma = \beta$, $\mu_2 = g_1$, we find no oscillations when $\beta \leq \frac{2}{3}$.

##### S1.2.3 Interpretation

Notice from the equilibrium solution (S23) that there are three singular values for the functional response exponents

- $\beta = 1$: predator abundance does not depend on predator mortality $\mu_2$ (the predator is strictly determined by its resource's productivity)

- $\gamma = 1$: prey abundance does not depend on prey growth $g_1$ (the prey is strictly determined by its predator)
- $\beta + \gamma = 1$: the exponent goes to infinity, meaning that there is no solution without self-regulation

When we look at how abundances vary with the parameters, we can discern two regimes:

- $\beta + \gamma > 1$ (including Lotka-Volterra $\beta = \gamma = 1$) gives that prey and predator abundance *decrease* with $A_{12}, \varepsilon$ and *increase* with $g_1$, $\mu_2$
- $\beta + \gamma < 1$ (which we showed to be unstable) gives the opposite; in particular, all abundances *decrease* with $g_1$

Some comments:

- For $\beta + \gamma > 1$, it may seem surprising that predator equilibrium abundance *increases* with its mortality $\mu_2$ if there is saturation $\beta < 1$. Intuitively, this is because increasing $\mu_2$ means higher predator mortality every year, but also allows much more prey to persist, leading to more growth. So we have an equilibrium with higher overall biomass and higher turnover (more predators die and are born each year).
- Still for $\beta + \gamma > 1$, equilibrium predator abundance *decreases* with $A_{12}$ and with $\beta + \gamma$ meaning that if a predator can control their predation behavior, they should strive to attack less in order to achieve larger populations. See however Sec. S2 for an explanation of why this might not happen in an adaptive context.
- By contrast, we saw that this solution is always unstable[2] for $\beta + \gamma < 1$. This means that abundances will either crash, or grow until self-regulation cannot be neglected. The risk of a crash (an emergent Allee effect) is only present if the unstable value of $B_2^*$ is larger than the extinction threshold 1, which requires large attack rates[3] $\tilde{A} > \tilde{A}_{\max}$.

**To summarize**

- $\beta + \gamma > 1$: a stable equilibrium without self-regulation exists, where the predator survives only for low attack rates $\tilde{A} < \tilde{A}_{\max}$,
- $\beta + \gamma < 1$:
  - for low attack rates $\tilde{A} < \tilde{A}_{\max}$, predator and prey abundance will grow until at least one of them is limited by self-regulation
  - for large attack rates $\tilde{A} > \tilde{A}_{\max}$, the predator-prey pair experiences a sort of Allee effect (and may either crash or grow until limited by self-regulation)

[2] This instability is hinted at by the fact that all abundances decrease with $g_1$: in a stable equilbrium, there should be positive bottom-up consequences of increasing basal productivity.

[3] We recall $\tilde{A} = A_{12}\varepsilon/\mu_2$, see Sec. S1.2.1.

- overall, without self-regulation, equilibrium abundances *increase* if we *decrease* attack rate and exponents $\beta, \gamma$.

#### S1.3 Self-regulated equilibria

##### S1.3.1 Resource self-regulation

If we assume that resources are limited by self-regulation, while predators are not ($B_2 \ll \mu_2/D_2$), we find the much more intuitive regime of donor control,

$$0 \approx g_1 B_1 - D_1 B_1^2 \tag{S32}$$

$$0 \approx -\mu_2 B_2 + \varepsilon A_{12} B_1^\beta B_2^\gamma \tag{S33}$$

with solution

$$B_1^* = \frac{g_1}{D_1} \tag{S34}$$

$$B_2^* = \left(\frac{\varepsilon A_{12}}{\mu_2}\right)^{\frac{1}{1-\gamma}} \left(\frac{g_1}{D_1}\right)^{\frac{\beta}{1-\gamma}} \tag{S35}$$

Since $\gamma \leq 1$, we see here that predator abundance $B_2^*$ always decreases with mortality $\mu_2$, increases with attack rate $A_{12}$ and with resource biomass $g_1/D_1$, and is larger than 1 (the extinction threshold) if

$$\tilde{A} > \tilde{A}_{\min} \equiv \left(\frac{D_1}{g_1}\right)^\beta \tag{S36}$$

where we recall our definition for the rescaled attack rate $\tilde{A} = A_{12}\varepsilon/\mu_2$.

##### S1.3.2 Consumer self-regulation

If $B_2 \gg \mu_2/D_2$ but $B_1 \ll g_1/D_1$, then only consumer self-regulation applies and we have

$$0 \approx g_1 B_1 - A_{12} B_1^\beta B_2^\gamma \tag{S37}$$

$$0 \approx -D_2 B_2^2 + \varepsilon A_{12} B_1^\beta B_2^\gamma \tag{S38}$$

which has the solution

$$B_2^* = \left(\frac{D_2}{\varepsilon A_{12}}\right)^{\frac{1}{2\beta+\gamma-2}} \left(\frac{\varepsilon g_1}{D_2}\right)^{\frac{\beta}{2\beta+\gamma-2}} \tag{S39}$$

We see that this equilibrium behaves like the no-self-regulation equilibrium, but with a singular line of exponents at $2\beta + \gamma = 2$ (e.g. $\beta = \gamma = 2/3$) rather than $\beta + \gamma = 1$ (e.g. $\beta = \gamma = 1/2$).

Hence, consumer self-regulation entails a less fundamental change of behavior than resource self-regulation, but simply shifts all our conclusions above. We can intuitively see how this behavior generalizes if we assume losses $L_2 = l_2 B_2^\delta$ with an arbitrary exponent $\delta > \beta + \gamma$.

##### S1.4 Synthesis

We now discuss the general picture over all equilibria.

- If $\beta + \gamma > 1$, the no-self-regulation equilibrium is stable. If $\beta + \gamma < 1$, it is unstable, and abundances will either crash or grow until self-regulation becomes important.
- In the no-self-regulation equilibrium, larger predator abundances are obtained by *decreasing* attack rate $A_{12}$ and exponents $\beta + \gamma$
- If $\beta + \gamma < 1$, populations either crash or reach the equilibrium where resource self-regulation dominates. In that equilibrium, larger predator abundances are obtained by *increasing* attack rate and exponents
- Hence, predator populations with $\beta + \gamma$ close to 1, for instance $\beta, \gamma \approx 1/2$, will typically be the largest stable populations.
- This requires both the self-regulated and non-self-regulated equilibria to allow persistence of the predator, i.e. the attack rate[4] must fall in the range
$$\left(\frac{D_1}{g_1}\right)^\beta = \tilde{A}_{\min} < \tilde{A} < \tilde{A}_{\max} = \left(\frac{g_1 \varepsilon}{\mu_2}\right)^\beta \tag{S40}$$
We notice that this range of persistence expands with $g_1$, and shrinks when $\beta \to 0$, hence saturation cannot be too strong.

#### S2 Adaptive dynamics

##### S2.1 Context

In the main text, we discuss the potential for an adaptive explanation of the observed exponents. If self-regulation $D_1, D_2$ is sufficiently weak to allow the non-self-regulated equilibrium when $\beta + \gamma > 1$, then predator abundance can be maximized by bringing $\beta + \gamma$ close to 1 (and retaining moderate attack rates).

We now consider under which circumstances a species could indeed adapt to maximize its abundance by consuming less of the resource.

Our basic intuition is that larger populations of "slower" consumers can prevail if resource depletion is not the only source of competition, but species at the same trophic level also experience strong direct competition (through aggression, or pathogens) regardless of resource levels. This is easy to see when considering two distinct species, e.g. lions displaying aggressive behavior towards hyenas. If direct inter-species competition is large compared to intra-species competition, an abundant consumer can prevent its competitor from invading in smaller populations, even if the competitor would be faster and better at exploiting the resource.

[4] We recall $\tilde{A} = A_{12}\varepsilon/\mu_2$, see Sec. S1.2.1.

A thornier question arises when we do not consider two distinct species, but mutants within the same species that develop slightly faster consumption (e.g. higher attack rate $A_{12}$ or exponents $\beta, \gamma$). In that case, inter-phenotype competition cannot be much larger than intra-phenotype competition, since they are part of the same species, and the aggressive behaviors or pathogens of non-mutants *a priori* cannot choose to target mutants dramatically more than non-mutants. Since faster consumption provides an instantaneous advantage in growth rate, we expect these mutants to invade and possibly replace the original population.

We will show that, indeed, faster consumers prevail in a basic model of adaptive dynamics, leading to maladaptive evolution: attack rates will increase, and populations decrease, until extinction.

But we will then show that this maladaptive evolution can be stopped if direct competition exists and increases with attack rates (independently of resource levels) – for instance, if individuals that search for or capture more food are met with more direct aggression from their conspecifics. This could be a basis for the widespread interference and territorial behaviors of consumers, displayed most spectacularly by higher vertebrates.

#### S2.2 Maladaptive evolution

A simple way to model adaptive dynamics is to assume that in one evolutionary step, a single individual mutates and obtains different values of $\beta, \gamma, A_{12}$. We then see whether this new phenotype (whose abundance we denote by $b_2$, by opposition to $B_2$ the resident population) can invade from an initial abundance $b_2 = 1$. At the end of the ecological dynamics, we choose either the resident or mutant phenotype to survive into the next evolutionary step, with probabilities proportional to their final abundances.

We thus write a new set of equations including this new phenotype. One important difficulty is that, since the functional response is nonlinear, we cannot simply add the contributions of both phenotypes. Since it is non mechanistic, there is also no grounded way of deriving the correct expression for their contributions.

We can however make a purely qualitative statement that does not depend on our precise model. Before the mutation, we can arbitrarily split the predator population in two subcomponents of sizes $B_2 = B_2^* - 1$ and $b_2 = 1$ with the same parameters, which must both be at equilibrium simultaneously.

$$\frac{dB_2}{dt} = \phi_2(B_1, B_2, b_2) = -\mu_2 B_2 - D_2 B_2 (B_2 + b_2) + F(B_1, B_2, b_2) = 0$$
$$\frac{db_2}{dt} = \phi_2'(B_1, b_2, B_2) = -\mu_2 b_2 - D_2 b_2 (B_2 + b_2) + F(B_1, b_2, B_2) = 0 \quad \text{(S41)}$$

To know whether $b_2$ will invade after a mutation, say an increase of attack rate $A_{12} \to A_{12} + \delta A$, we must simply know whether

$$\frac{\partial \phi_2'}{\partial A_{12}}(B_1 = B_1^*, B_2 = B_2^* - 1, b_2 = 1) > 0 \quad \text{(S42)}$$

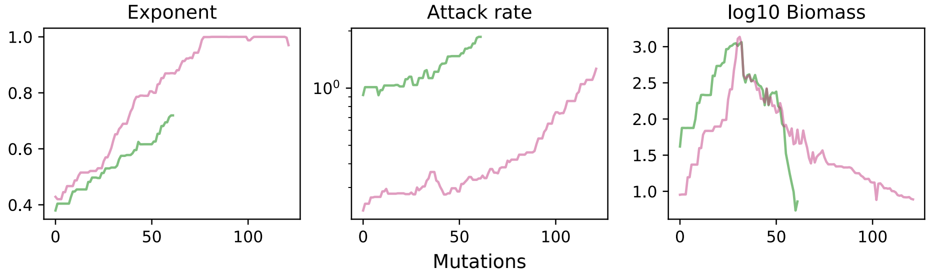

Figure S2: Maladaptive evolution in adaptive dynamics simulations. Each mutation happens in 1 predator individual within an equilibrium resident population of predators. This individual acquires slightly shifted attack rate $A_{12}$ and exponents $\beta$ and $\gamma$ (here we maintain $\beta = \gamma$ and shift both simultaneously). We then run ecological dynamics and the phenotype with the largest population at the end of the run is selected as the resident population for the next mutation. We repeat this process over 200 mutations or until extinction. This evolutionary process can first lead to an increase, but eventually a decrease of the standing predator biomass, down to extinction, as faster consumers (larger attack rate and exponents) outcompete slower consumers but fail in the long term, driving both extinct.

i.e. whether a population starting at equilibrium and suddenly shifting to a larger attack rate will increase at first (before resource depletion may bring it down). Since $A_{12}$ appears (so far) only as a constant inside $F$, it is unavoidable that a larger attack rate will lead to an initial growth of the mutant phenotype. This will drive the resource down in the long term, and classic competition theory suggests that the faster consumer will prevail.

As a consequence, mutations will drive an increase in attack rates and a decrease in populations all the way to collapse, as we show in simulations in Fig. S2. These simulations follow the description above, but for simplicity, we assume

$$\begin{aligned}
\frac{dB_1}{dt} &= \phi_1(B_1, B_2, b_2) \approx g_1 B_1 - D_1 B_1^2 - A_{12} B_1^{\beta} B_2^{\gamma} - A'_{12} B_1^{\beta'} b_2^{\gamma'} \\
\frac{dB_2}{dt} &= \phi_2(B_1, B_2, b_2) = -\mu_2 B_2 - D_2 B_2 (B_2 + b_2) + \varepsilon A_{12} B_1^{\beta} B_2^{\gamma} \\
\frac{db_2}{dt} &= \phi'_2(B_1, B_2, b_2) = -\mu_2 b_2 - D'_2 b_2 (B_2 + b_2) + \varepsilon A'_{12} B_1^{\beta'} b_2^{\gamma'} \qquad \text{(S43)}
\end{aligned}$$

where $D'_2 = D_2$, which is compatible with our scaling only when either $B_2$ or $b_2$ is nonzero but not both. We do not expect more sophisticated decompositions of the nonlinear functional response to induce a qualitatively different behavior.

##### S2.3 Surviving through direct competition

The crucial point above is that the competition coefficient $D_2' = D_2$ is the same for inter- and intra-phenotype competition. We cannot have an independent competition term arising from $B_2$ in the equation of $b_2$, contrary to competition between two different species, because we assume that individuals of the resident population cannot intrinsically recognize mutants and punish them.

However, if competition depends on the *traits* of the phenotypes, then it can prevent malaptive evolution, provided the resident population is abundant enough. In the simplistic model (S43) above, if we assume for instance that competition increases proportionally with attack rate and exponents

$$D_2 = A_{12}\beta\gamma d_2, \qquad D_2' = A_{12}'\beta'\gamma' d_2 \tag{S44}$$

with some constant $d_2$, then for large enough resident population $B_2^*$ we can have

$$\frac{\partial \phi_2'}{\partial A_{12}'}(B_1^*, B_2^*, b_2 = 1),\ \frac{\partial \phi_2'}{\partial \beta'}(B_1^*, B_2^*, b_2 = 1),\ \frac{\partial \phi_2'}{\partial \gamma'}(B_1^*, B_2^*, b_2 = 1) < 0 \tag{S45}$$

which ensures that the resident population cannot be invaded. In the simplified equations (S43), we find for instance $\partial\phi_2'/\partial A_{12}' < 0$ if the resident population exceeds a threshold

$$B_2^* > \frac{\varepsilon B_1^{\beta'}}{\beta'\gamma' d_2} \tag{S46}$$

and similar conditions can be found for derivatives with respect to $\beta'$ and $\gamma'$. It is crucial that this competition term not be tied to *success* at finding resources (i.e. be dependent on prey density), since it would then vanish if the mutant can deplete the resource to lower levels than the resident population can survive on. Instead, it must target individuals who *search* for resources more, i.e. who intrinsically have larger $A_{12}$, $\beta$ and $\gamma$. This feature seems plausible in particular for territorial and aggressive behavior.

#### S3 Allometric scaling

##### S3.1 Biomass-production and predator-prey scalings

Let us assume a food chain where each level has biomass $B_i$, production $P_i$, and some internal losses (due to mortality, self-regulation, etc) that scale like $B_i^\delta$ with some exponent $\delta$

$$\frac{dB_i}{dt} = -\underbrace{\mu B_i^\delta}_{\text{internal losses}} + P_i - \frac{1}{\varepsilon}P_{i+1} \tag{S47}$$

with $\varepsilon$ the trophic efficiency. In general, production $P_i$ should depend on both $B_i$ and $B_{i-1}$. For instance, in Lotka-Volterra models, $P_i \sim B_i B_{i-1}$.

If we ignore losses $\mu = 0$, then at equilibrium $\varepsilon P_i = P_{i+1}$. With Lotka-Volterra, we can simplify $B_i$ on both sides and get $\varepsilon B_{i-1} \sim B_{i+1}$. In other words, a Lotka-Volterra chain without losses does *not* give a simple scaling relationship between adjacent levels $B_i$ and $B_{i-1}$, or between production and biomass at the same level $P_i$ and $B_i$. It imposes a relationship between levels that are *two* steps apart, $B_{i-1}$ and $B_{i+1}$.

There are two possibilities explaining an allometric scaling between $P_i$ and $B_i$:

- either this scaling is imposed by some non-Lotka-Volterra functional response itself, i.e. we simply assume in the model that $P_i(B_i, B_{i-1}) \sim B_i^x$ (which we will see later with Arditi-Ginzburg style ratio-dependent functional response).
- or the dynamics impose that scaling via the equilibrium condition $dB_i/dt = 0$.

To ensure a scaling that results from the dynamics, we can assume that internal losses are comparable to, or larger than, predation losses

$$\mu B_i^\delta \gtrsim \frac{1}{\varepsilon} P_{i+1}. \tag{S48}$$

In that case, biomass will grow until the equilibrium relation

$$P_i \sim \mu B_i^\delta \tag{S49}$$

between growth and losses is reached. Thus, a scaling relationship emerges at equilibrium that is not imposed out of equilibrium (contrary to assuming that this scaling relationship is inside the functional response, in which case it holds at all times). The condition for indeed observing empirically $P_i \sim B_i^\delta$ is that the prefactor $\mu$ should not vary significantly, or at least should not be a main driver of the variation of biomass $B_i$ between ecosystems; otherwise, $\mu$ and $B_i$ will be correlated in some ways that can alter the observed relationship (for instance, if we see a correlation $\mu \sim 1/B_i$ across distinct ecosystems, we will observe $P_i \sim B_i^{\delta-1}$ instead).

##### S3.1.1 Functional response

The usual definition of the functional response is $f$ such that

$$P_i = B_i f(B_i, B_{i-1}) \tag{S50}$$

As in the previous appendix, let us assume that production has a power-law dependence in both levels (see below for examples)

$$P_i = \varepsilon a B_{i-1}^\beta B_i^\gamma \tag{S51}$$

with $a$ a measure of interaction strength (in the appropriate units given $\gamma$ and $\beta$).

##### S3.1.2 Scaling laws between adjacent levels

Then substituting (S49)

$$P_i = \varepsilon a \left(\frac{P_i}{\mu}\right)^{\gamma/\delta} \left(\frac{P_{i-1}}{\mu}\right)^{\beta/\delta} \tag{S52}$$

Hence we get the scaling relationship between the productions of adjacent levels

$$P_i = \xi P_{i-1}^{\alpha} \tag{S53}$$

with

$$\xi = \frac{\varepsilon a}{\mu^{(\gamma+\beta)/\delta}}, \qquad \alpha = \frac{\beta}{\delta - \gamma} \tag{S54}$$

Likewise, we get the scaling relationship between the biomasses of adjacent levels

$$B_i = \tilde{\xi} B_{i-1}^{\alpha} \tag{S55}$$

with

$$\tilde{\xi} = (\xi \mu^{\alpha-1})^{1/\delta} \tag{S56}$$

##### S3.1.3 Low predation loss condition

Our initial premise was that predation losses are similar or negligible compared to internal losses (S48). It is equivalent to

$$P_i \gtrsim \frac{1}{\varepsilon} P_{i+1} \tag{S57}$$

Now, this takes the form

$$\begin{aligned} P_i &\gtrsim \frac{\mu}{\varepsilon} P_i^{\alpha} \\ P_i &\gtrsim \left(\frac{\mu}{\varepsilon}\right)^{1/(1-\alpha)} \qquad \forall i \end{aligned} \tag{S58}$$

(if $1 > \alpha > 0$ which seems reasonable). Since production must always be smaller at higher levels, this relation cannot in general hold at every level for infinitely long chains (unless $\alpha = 1$ and $\mu < \varepsilon$), and requires sufficiently high primary productivity.

##### S3.1.4 Summary

So far we have obtained two different scalings:

$$P_i = r B_i^{\delta}, \qquad B_{i+1} = \tilde{\xi} B_i^{\alpha}$$

with $\alpha = \beta/(\delta - \gamma)$ and coefficients detailed above.

#### S3.2 Lotka-Volterra biomass pyramid

Assuming a Lotka-Volterra functional response, we get

$$P_i \sim B_i B_{i-1}, \qquad \beta = \gamma = 1 \tag{S59}$$

Then a simple biomass pyramid

$$B_i \sim B_{i-1}, \qquad \alpha = 1 \tag{S60}$$

requires $\delta = 2$. The only choice is to have quadratic losses in our dynamical equation (usual self-competition), and then we get

$$P_i \sim B_i^2. \tag{S61}$$

So we don't have the same scaling for production versus biomass ($\delta = 2$) and for predators versus prey ($\alpha = 1$).

#### S3.3 Same scaling for production and predators

Data discussed in Hatton et al. [3] seems to indicate

$$\delta \sim \alpha = \frac{\beta}{\delta - \gamma} \tag{S62}$$

which boils down to the condition

$$\delta(\delta - \gamma) = \beta \tag{S63}$$

$$\delta^{\pm} = \frac{1}{2}(\gamma \pm \sqrt{\gamma^2 + 4\beta}) \tag{S64}$$

Hereafter we only consider the + solution which is positive, and we treat some simple subcases.

##### S3.3.1 Lotka-Volterra functional response

Again assuming Lotka-Volterra functional response

$$P_i \sim B_i B_{i-1}, \qquad \beta = \gamma = 1 \tag{S65}$$

Then

$$\delta = \frac{1 + \sqrt{5}}{2}, \qquad P_i \sim B_i^{1.618...} \tag{S66}$$

(this is the golden ratio). To get $P_i$ that increases *sub*linearly with $B_i$, we will need a functional response that increases more slowly with $B_i$ and $B_{i-1}$.

###### S3.3.2 Linear biomass-production scaling

To get a linear scaling relationship $P_i \sim B_i$ ($\delta = 1$), we need

$$1 - \gamma = \beta \tag{S67}$$

for instance $\gamma = \beta = 1/2$. More generally, it means that the production can be written as

$$P_i = \varepsilon a B_i^{\gamma} B_{i-1}^{1-\gamma} \tag{S68}$$

$$= B_i f(B_i, B_{i-1}) \tag{S69}$$

with a ratio-dependent functional response[4]

$$f(B_i, B_{i-1}) = \varepsilon a \left(\frac{B_i}{B_{i-1}}\right)^{\gamma-1}. \tag{S70}$$

For instance, we can obtain this in some limits from the functional reponse

$$P_i = B_i f(B_i, B_{i-1}) \propto \frac{B_i B_{i-1}}{1 + IB_i + HB_{i-1}}. \tag{S71}$$

with $I$ predator interference, and $H$ handling (1/half saturation) in a Type-2 functional response.

The limit $I \to \infty$ gives $\gamma = 0$, $\beta = 1$ (only the number of prey counts). The limit $H \to \infty$ gives $\gamma = 1$, $\beta = 0$ (only the number of predators counts).

###### S3.3.3 Empirical allometric laws

To get sublinear scaling $\delta \approx 0.75$ as in Hatton et al.[3], we need

$$\frac{3}{4}(\frac{3}{4} - \gamma) = \beta \tag{S72}$$

For example for $\gamma = \beta$ we get

$$\gamma = 9/28 \approx 0.32 \tag{S73}$$

Hence we have approximately

$$P_i \propto (B_i B_{i-1})^{0.32}, \tag{S74}$$

Various exponents can emerge from spatial agregation[5;6], but further explanation would be needed for the robustness of this specific exponent.

### S4 Empirical analysis

#### S4.1 Data description

The data comprises 62 tab-separated lines with the following fields:

- `source`: reference, see bibliography below
- `c12`: kill rate (prey/predator/year)
- `study area`: extension of the study area (km$^2$)

and for each species $i = 1, 2$:

- `species`: species name
- `w`: body mass (kg)
- `m`: mass-specific metabolic rate (W/kg)
- `density`: numeric density over the area of study (individuals/km$^2$)

All properties except `source`, `species` and `study area` are further broken down into multiple columns:

- `used`: value used in our analysis, converted in the above units
- `source`: value in the reference
- `source units`: units in the reference
- `type`: whether the source value was directly given in the text (number), taken as an average over such numbers (`average data`) or a visual average from a figure (`average data from figure`), obtained by adding multiple numbers (`addition`), or derived from allometric scaling between mass and metabolism (`alloRelationsMW`)
- `notes`: other informations relevant to how this value was obtained, e.g. external sources used, rationale behind a conversion.

#### S4.2 Data collection

We collected 32 studies (see dataset attached to the Supplementary Materials) reporting field kills $K$ over an extended period of time (typically a year or more). We show in Fig. S3 the scalings relating $K$ to biomass densities $B_1$, $B_2$ and mass-specific metabolic rates $m_1$ and $m_2$, combined into "energy" density $E_i = m_i B_i$.

We focused on higher terrestrial vertebrates, due to the possibility of direct observation of predation in the form of kills, carcasses, nest visits, etc. This choice likely introduces a taxonomic bias which will need to be ascertained by contrasting our results with other sources of data, notably feeding experiments on invertebrates, and (often more indirect) measurements of predation in aquatic systems. Even within terrestrial vertebrates, we found a highly uneven number of studies depending on predator species, with some charismatic species and top predators (e.g. wolf, lion) drawing much more attention. Rather than attempt an exhaustive or balanced sampling of data over predator-prey pairs,

| | $\beta$ | $\gamma$ | $\nu$ |
|---|---|---|---|
| Aggregated (N=32) | $0.47 \pm 0.17$ | $0.30 \pm 0.18$ | $0.43 \pm 0.28$ |
| Disaggregated (N=62) | $0.54 \pm 0.12$ | $0.23 \pm 0.12$ | $0.19 \pm 0.18$ |

Table S1: Best-fit exponents $\beta$ and $\gamma$ and their standard deviations (computed by bootstrapping, very similar to direct estimates from the linear fit) for data aggregated by study and by species, or fully disaggregated, showing robustness against uneven resolution.

we designed a simple test to check whether the uneven resolution of this sampling could significantly alter our conclusions.

A number of studies contained multiple sites, multiple temporal measurements, or multiple predator-prey species pairs. For some such studies, we collected all possible measurements as distinct data points; for others, we only took a single measurement or an average or sum, in such a way that all choices occur at least once in the dataset. We then tested two options: retaining all 62 datapoints as if they were independent, or aggregating results within each of the 32 studies (taking an average over sites and time points and summing over prey species for each predator species).

As we show in Table S1, these options did not cause a significant change in any of the exponents in the fitted power-laws (S75) and (S76) described below. We note that the exponent $\nu$ in (S76) is closer to 0 when data is disaggregated, while close to 0.5 when aggregated. This exponent contributes little to the fit quality, see Table 1 in the main text. By contrast, the scaling exponents $\beta$ and $\gamma$ seem robust to over- or under-representing certain subclasses within the data, suggesting that they are truly meaningful.

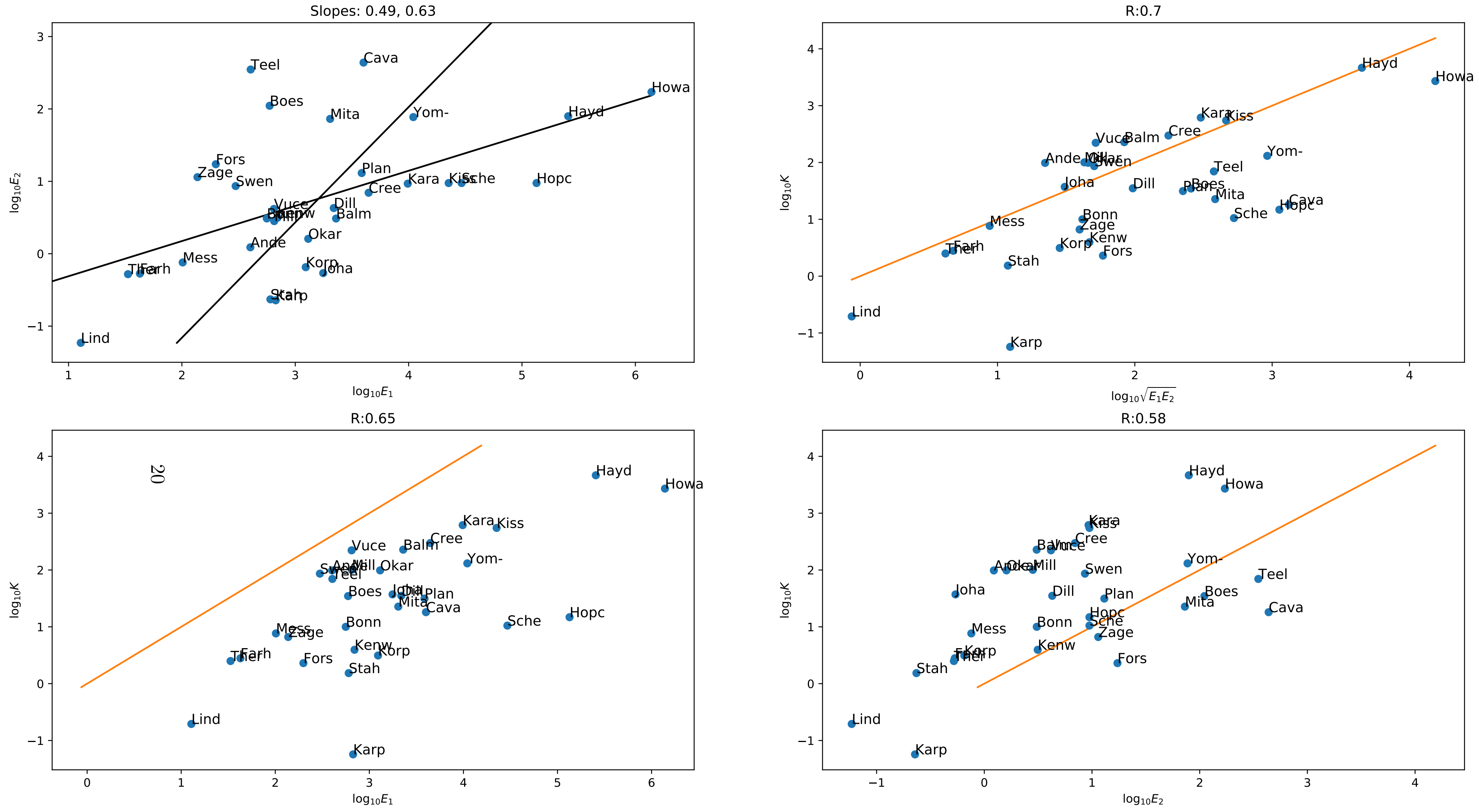

Figure S3: Empirical relationship between predation $K$ and energy densities $E_1 = m_1B_1$, $E_2 = m_2B_2$. Top-left: $E_2$ versus $E_1$; solid lines represent fits $E_1 \sim E_2^{0.49}$, $E_2 \sim E_1^{0.63}$. Top-right: $K$ versus $\sqrt{E_1E_2}$. Bottom-left: $K$ versus $E_1$. Bottom-right: $K$ versus $E_2$. Correlation coefficients are shown in panel headings, and solid lines denote the 1:1 relationship. Each point is labelled by the first four letters of the reference (data have been aggregated within a study).

#### S4.3 Comparison of functional responses

As noted in the main text, we consider two chief candidates given the possibility of both saturation and interference in our data: the power-law expression

$$K_i \equiv K(B_i, B_{i+1}) = A_{i,i+1} B_i^\beta B_{i+1}^\gamma \tag{S75}$$

and its ratio-dependent version where

$$K = A m_2 B_2 \left(\frac{m_1}{m_2}\right)^\nu \left(\frac{B_1}{B_2}\right)^\beta \tag{S76}$$

on one hand, and the DeAngelis-Beddington model [7;8])

$$K(B_i, B_{i+1}) = \frac{A B_i B_{i+1}}{1 + H B_i + I B_{i+1}} \tag{S77}$$

on the other.

By performing a fit of these two models against $B_1$ and $B_2$ in log-log space, we find that the rational (DeAngelis-Beddington or DAB) model is out of its range of relevance: the fitted coefficients $H$ and $I$ give

$$H B_1 + I B_2 \gg 1 \tag{S78}$$

for almost all data points $B_1$,$B_2$ (in 97% of cases, the saturation threshold is exceeded by a factor of 100). This means that the model only applies in the completely saturated regime, where we recover either donor control (if $IB_2$ is larger) or consumer control (if $HB_1$ is larger).

In its saturated regime, the DAB model can, peculiarly, approximate our power-law result due to the colinearity of $B_1$ and $B_2$. Indeed, if we assume

$$\log B_2 = \log B_1 + b + \xi \tag{S79}$$

with $a$ and $b$ two constants, and $\xi$ a relatively small random effect term that varies between systems, we then have

$$\log \sqrt{B_1 B_2} = \frac{1}{2}\left(2 \log B_1 + b + \xi\right) \tag{S80}$$

$$= \log B_1 + \frac{b+\xi}{2} \tag{S81}$$

But we can also see that, if $e^b = H/I$,

$$\begin{aligned}
\log(1 + HB_1 + IB_2) &= \log\left(1 + B_1\left(H + I e^{b+\xi}\right)\right) \\
&\approx \log B_1 + \log H + \log(1 + e^\xi) \\
&\approx \log B_1 + \log H + \log(2 + \xi) \\
&= \log B_1 + \log H + \log 2 + \log\left(1 + \frac{\xi}{2}\right) \\
&\approx \log B_1 + \log H + \log 2 + \frac{\xi}{2} \\
&= \log B_1 + \frac{b+\xi}{2} \qquad \text{if} \log H + \log 2 = \frac{b}{2}
\end{aligned} \tag{S82}$$

where we use $\log(1+x) \approx \log x$ (for $x \gg 1$) and $e^x \approx 1+x$ , $\log(1+x) \approx x$ (for $x \ll 1$). Hence the two results (S81) and (S82) are identical to first order in $\xi$ provided that saturation and interference are adjusted precisely so that

$$H = \frac{1}{2}e^{b/2}, \qquad I = \frac{1}{2}e^{-b/2}. \tag{S83}$$
