## Supplementary material for "A macro-ecological approach to predators’ functional response": Simulation code: simu_suppmats.py

```
import numpy as np, matplotlib.pyplot as plt, pandas as pd
import sys, os
from matplotlib import cm


"""
README:
This is the code associated with 'A macro-ecological approach to predators' functional response' M.Barbier et al. bioRxiv 832220 (2020)

This code should be run with Python 2.7 (works with package versions: NumPy 1.15.4, Matplotlib 2.2.2, Pandas 0.23.0)

When it is run, two functions will be executed one after the other:
theo() will first return a single plot containing all the theoretical results on the simple model analyzed in the article
twopred(loop=True) will then simulate adaptive dynamics (see details in Supplementary Appendices) and return the corresponding plot

"""


def ifelse(x,a,b):
    if x:
        return a
    return b
    
########################################## SIMULATIONS ###################################
    
def onerun(tmax=200.,dt=0.01,a=0.7,b=0.7,g1=1.,r2=2.,A=1.,eps=0.8,D1=0.1,D2=0,death=10**-8,print_msg=1,**dic):
    r2stab=g1*(1-a)/(1-b)
    if r2 is None:
        r2=r2stab*2
    B=np.ones(2)
    r2e=float(r2)/eps
    ## SCENARIO 1: NO SELF REG
    if (a+b)>1:
        eq1=(np.array([g1**(1-b) * r2e**b,g1**a * r2e**(1-a)])/A)**(1/(a+b-1) )
    else:
        eq1=np.array((np.inf,np.inf))
    ## SCENARIO 2: RESOURCE SELF REG
    eq2=np.array([g1/D1, (  A/r2e * (g1/D1 )**a )**(1./(1.-b)) ]  )
    if eq1[0]/eq2[0] < 2 and eq2[0]/eq1[0]<2:
        # Same strength of self-reg and other-reg
        eq=np.array([g1/D1/2., (  A/r2e * (g1/D1/2. )**a )**(1./(1.-b)) ]  )    
    elif eq1[0] > eq2[0]:
        eq=eq2
    else:
        eq=eq1
    B=eq+np.ones(2)*0.01
    if 'x0' in dic:
        B=dic['x0']
    def dBdt(t,B):
        B1,B2=np.clip(B,0,None)
        eq1=g1*B1 - A*B1**a * B2 ** b - D1*B1**2
        eq2=-r2*B2 + eps*A*B1**a * B2 ** b - D2*B2**2
        return np.array([eq1,eq2])

    tab=[]
    t=0
    print B
    while t< tmax:
        if (t%10. < dt) and print_msg:
            print t
        diff=dBdt(t,B)
        diff[B<death]=np.clip(diff[B<death],0,None)
        B=np.clip(B+dt*diff,0,None)
        t+=dt
        B1,B2=B
        if B1< death and 0:
            print '     DEATH', g1*B1, r2*B2, A*B1**a * B2 ** b
            B2=death/2
        tab.append({'t':t,'B1':B1,'B2':B2, 'B1eq':eq[0], 'B2eq':eq[1]})
        if np.max((B1,B2))< death:
            break
    tab=pd.DataFrame(tab)
    return tab

    
 
def twopred(path='./',fname='simuloop.csv',loop=0,death=1.,deltab=0.03,deltaA=0.1,r=1,D=0.0,C=1,nu=1, **kwargs):
    #competition between two predators
    def run(tmax=1000,maxdt=1,a=0.99,b=0.99,g1=1,r2=0.01,A=1,eps=0.8,D1=0.01, r3=0.01,A3=1.,a3=1,b3=1, D2=1,D3=1,C23=1,C32=1,nu2=0,nu3=0,**kwargs):
        r2e=r2/eps
        r3e=r3/eps
        
        ## EQUILIBRIA
        if (a+b)>1:
            Beq1=(np.array([g1**(1-b) * r2e**b,g1**a * r2e**(1-a)])/A)**(1/(a+b-1) )
        else:
            Beq1=np.array((np.inf,np.inf))
        ## SCENARIO 2: RESOURCE SELF REG
        Beq2=np.array([g1/D1, (  A/r2e * (g1/D1 )**a )**(1./(1.-b)) ]  )
        if Beq1[0] > Beq2[0]:
            Beq=Beq2
        else:
            Beq=Beq1
            

        
        B=np.array([max(g1/D1,death*100),death*20,death*10])
        if 'x0' in kwargs:
            B=np.array(kwargs['x0'])
        if kwargs.get('avoid_allee',1):
            B[0]=np.max([ B[0],2* (r2/eps *B[1]**(1-b) )**(1./a) ,2* (r3/eps *B[2]**(1-b3) )**(1./a3)   ])
        #print B
        def dBdt(t,B):
            B1,B2,B3=np.clip(B,0,None)
            eq1=g1*B1 - A*B1**a * B2 ** b - A3*B1**a3 * B3**b3 - D1*B1**2
            eq2=-r2*B2 + eps*A*B1**a * B2 ** b - D2  * A**nu3 * B2*(B2   +C23* B3  )
            eq3=-r3*B3 + eps*A3*B1**a3 * B3 ** b3  - D3 *A3**nu2 * B3*(C32*B2  +B3)
            return np.array([eq1,eq2,eq3])

        #print '         eq:', dBdt(0,B), (np.array([g1**(1-b) * r2e**b,g1**a * r2e**(1-a)])/A)**(1/(a+b-1) )
        tab=[]
        t=0
        while t< tmax and np.max(B)>death:
            
            diff=dBdt(t,B)
            diff[B<death]=np.clip(diff[B<death],10**-15,None)
            diff[B==0]=0
            dt= min(maxdt,min(tmax-t,np.min(np.abs(B/diff)[B>death] )*0.05))
            if (t%(tmax/10) < dt) and kwargs.get('print_msg',1):
                print t, B
            B=np.clip(B+dt*diff,0,None)
            t+=dt
            B1,B2,B3=B
            tab.append({'t':t,'B1':B1,'B2':B2,'B3':B3,'eqB2':Beq[1], 'eqB1': Beq[0], })
        tab=pd.DataFrame(tab)
        return tab

    
    dic={'g1':10, 'a':0.8,'b':0.8, 'A':0.01, 'D1':0.01,'D2':D,'D3':D,'r2':r,'r3':r,'C23':C,'C32':C,'eps':.8, 'nu2':nu,'nu3':nu}
    dic['x0']=[max(dic['g1']/dic['D1'],death*100),max(death*10,dic['g1']/dic['A']) , 0]
    dic.update(kwargs)
    
    if loop:
        ntrials=kwargs.get('ntrials',5)
        ngen=kwargs.get('ngen',500)
        if 'rerun' in sys.argv or not fname in os.listdir(path):
            table=[]
            for trial in range(ntrials):
                print '=================== TRIAL', trial
                beta=np.random.uniform(0.,1)
                A=10**(np.random.uniform(-2,0))
                dA=db=0
                for gen in range(ngen):
                    dic['a']=dic['b']=beta
                    dic['a3']=dic['b3']=np.clip(beta+np.random.uniform(-1,1)*deltab,0.,1)
                    dic['A']=A
                    dic['A3']=A*(1+np.random.uniform(-1,1)*deltaA)

                    df=run(print_msg=0,**dic)
                    thresh=int(df.shape[0]*3/4)
                    B1,B2,B3=np.mean(df[['B1','B2','B3']].values[thresh:],axis=0)
                    if gen%10==0 or 'analyze' in sys.argv:
                        print 'Generation',gen, B2,B3,A,beta
                    if max(B2,B3)<=death or B1<=death:
                        break

                    oldA,oldbeta=A,beta
                    if B2>B3:#np.random.random()< np.exp( 1*(B2-B3)/(B2+B3) ) :
                        A,beta,x0=dic['A'],dic['a'],B2
                    else:
                        A,beta,x0=dic['A3'],dic['a3'],B3
                    dA,db=(A-oldA)/oldA,beta-oldbeta


                    result={'gen':gen,'trial':trial,'beta':beta,'A':A,'logB':np.log10(x0),'logB1':np.log10(B1),
                        'dA':dic['A3']/dic['A']-1,'db':dic['a3']-dic['a'],
                        'accdA':dA,'accdb':db,'eqB1': df['eqB1'].values[-1],'eqB2': df['eqB2'].values[-1] }
                    table.append(result)
                    dic['x0']=(dic['g1']/dic['D1'],max(death*20,x0),max(death*10,x0/50.))
                    
                if 'analyze' in sys.argv:
                    dft=pd.DataFrame(table)
                    plt.subplot(221),plt.title('Attack rate {:.2g} (b) vs {:.2g} (r)'.format(dic['A'],dic['A3'])) #,plt.title('Exponent')
                    kw={}
                    plt.plot(df['t'],df['B1'],color='g',**kw)
                    plt.plot(df['t'],df['B2'],color='b',lw=2,**kw)
                    plt.plot(df['t'],df['B3'],color='r',lw=2,**kw)
                    plt.yscale('log')
                    plt.legend(['B1','B2','B3'])

                    plt.subplot(222)
                    plt.plot(dft['gen'],np.log10(dft['A']))
                    plt.plot(dft['gen'],dft['beta'])
                    plt.plot(dft['gen'],dft['logB'])
                    plt.plot(dft['gen'],dft['logB1'])
                    plt.legend(['log10 Attack rate','Exponent','log10 Biomass','log10 Ressource'])

                    plt.subplot(223),plt.xlabel('Exponent'),plt.ylabel('Attack rate')
                    plt.scatter(dft['db'],dft['dA'])
                    plt.scatter(dft['accdb'],dft['accdA'],marker='x')
                    plt.legend(['Proposed','Accepted'])
                    plt.subplot(224),plt.title('Power-law eq. B2')
                    plt.hist(dft['eqB2'])
                    plt.show()

                
            table=pd.DataFrame(table)
            table.to_csv(path+fname)
        else:
            table=pd.read_csv(path+fname)
        Bmin=np.min(table.groupby('trial')['logB'].mean())
        for trial, gp in table.groupby('trial'):
            lw=(1+np.mean(gp['logB']-Bmin) )
            color=cm.get_cmap('plasma')(gp['A'].mean() )
            plt.subplot(221),plt.title('Exponent')
            plt.plot(gp['gen'],gp['beta'],lw=lw,color=color,alpha=.5)
            plt.subplot(222),plt.title('Attack rate'),plt.yscale('log')
            plt.plot(gp['gen'],gp['A'],lw=lw,color=color,alpha=.5)
            plt.subplot(223),plt.title('Power-law eq'),plt.yscale('log')
            plt.plot(gp['gen'],gp['eqB2'],lw=lw,color=color,alpha=.5)
            plt.plot(gp['gen'],gp['eqB1'],lw=lw,color=color,alpha=.5,linestyle='--')
            plt.subplot(224),plt.title('log10 Biomass')
            plt.plot(gp['gen'],gp['logB'],lw=lw,color=color,alpha=.5)
    else:
        dic['a']=dic['b']=.75
        dic['A']=dic['A3']=0.03  
        dic['x0'][1]=0
        dic['x0'][2]=death*10
        dic['tmax']=50
        for beta,s in zip([.83,.95],[None,'--']):
            dic['a3']=dic['b3']=beta
            df=run(**dic)
            kw={'linestyle':s}
            plt.plot(df['t'],df['B1'],color='g',**kw)
            plt.plot(df['t'],df['B2'],color='b',lw=2,**kw)
            plt.plot(df['t'],df['B3'],color='r',lw=2,**kw)
            plt.plot(df['t'],np.ones(df.shape[0])*(dic['g1']**(1-dic['b3'])*(dic['r3']/dic['eps'])**(dic['b3'])/dic['A3'])**(1./(dic['a3']+dic['b3'] -1 )) ,color='g',lw=1,**kw)
            plt.plot(df['t'],np.ones(df.shape[0])*(dic['g1']**dic['a3']*(dic['r3']/dic['eps'])**(1-dic['a3'])/dic['A3'])**(1./(dic['a3']+dic['b3'] -1 )) ,color='r',lw=1,**kw)
            plt.yscale('log')
            plt.legend(['B1','B2','B3'])
    plt.show()


########################################## THEORY ###################################


def theo():
    ## MATRIX of beta,gamma

    dft={'eps':0.8,'D1':0.001,'D2':0.0, 'g1':1.,'r2':2. }

    def eqs(x, a=0, b=0, g1=0, r2=0, A=0, eps=0, D1=0, D2=0):
        B1, B2 = np.exp(np.clip(x, None, 20))
        eq1 = g1 - A * B1 ** (a - 1) * B2 ** b - D1 * B1
        eq2 = -r2 + eps * A * B1 ** a * B2 ** (b - 1) - D2 * B2
        return np.array((eq1, eq2))


    def Amin(D1=1,g1=1,r2=1,eps=1,a=1, **kw):
        return r2/eps* (D1/g1)**a

    def Amax(g1=1,r2=1,eps=1,a=1, **kw):
        return r2/eps* (g1*eps/r2)**a

    def geteq(a=0.7,b=0.7,g1=1.,r2=2.,A=1.,eps=0.8,D1=0.001,D2=0,shape=None,return_unstable=0,**kw):
        r2e=r2/eps
            
        if shape is None:
            shape=np.ones((1,1))
        else:
            shape=np.ones(shape)

        eq1=[ (x*shape /A)**(1./(a+b-1.))  for x in  [g1**(1-b) * r2e**b  ,g1**a * r2e**(1-a) ] ] 
        if return_unstable:
            return eq1
        eq1=np.array(eq1)
        eq1[0][np.where( (a+b)*shape <1 )]=np.inf
        eq1[1][np.where( (a+b)*shape <1 )]=np.inf
        
        eq2=[g1/D1*shape, (  A/r2e * (g1/D1 )**a )**(1./(1.-b)) *shape ]  
        B1,B2= np.minimum(eq1[0],eq2[0]),np.minimum(eq1[1],eq2[1])

        from scipy.optimize import root

        symbols=['B1','B2','a','b','g1','r2','A','eps','D1','D2']
        loc=locals()
        prms=pd.DataFrame({z:ifelse(hasattr(loc[z],'shape'),loc[z],
                                    loc[z]*np.ones(B1.shape) ).ravel()
                           for z in symbols })


        selfreg=(prms.D2*prms.B2 > prms.r2)
        prms.B2[selfreg]=np.minimum(prms.B2, (prms.B1**prms.a * prms.eps/prms.D2)**(1./(2-prms.b) )  )[selfreg]

        tab=[]
        for idx,row in prms.iterrows():
            args=(row.a, row.b,row.g1,row.r2,row.A,row.eps,row.D1,row.D2 )
            x0=np.array((row.B1,row.B2))
            class Tmp():
                success=False
            res,err=Tmp(),1

            methods=['hybr','lm',] #'krylov'
            while (err>10**-5 or not res.success) and methods:
                try:
                    res = root(eqs, x0=np.log(x0), args=args,method=methods.pop(0))
                except:
                    raise Exception('Error in rootfinder')
                err=np.max(np.abs(eqs(res.x,*args)))
            b1,b2=np.exp(res.x)
            if not res.success or err>10**-5:
                b1,b2=np.nan,0
            tab.append({'B1':b1,'B2':b2,'success':res.success,'err':err })

        tab=pd.DataFrame(tab)
        tab.B2[tab.success==False]=np.nan#B2.ravel()[tab.success==False]
        B1b,B2b=tab.B1.values.reshape(B1.shape),tab.B2.values.reshape(B1.shape)
        B1,B2=B1b,B2b

        return B1,B2,tab.err.values.reshape(B1.shape)


    def getstab(B1=0,B2=0,D1=0,D2=0,g1=0,r2=0,a=0,b=0,**kw):
        from scipy.linalg import solve_continuous_lyapunov as solve_lyapunov
        J12= a * (-g1 + D1*B1)
        J21= b * ( r2 + D2*B2)
        J11= -(2-a) * D1*B1 + (1-a)*g1
        J22= -(2-b) * D2*B2 - (1-b)*r2
        T=J11+J22
        D= J11 * J22 - J12*J21
        eig=(T+np.power(T**2-4*D+0j,.5 ) )/2.

        def getinv(J11,J12,J21,J22,B1,B2):
            if np.isnan([B1,B2]).any():
                return np.ones(4)*np.nan
            mat=np.array([[J11,J12],[J21,J22]])
            dempred=solve_lyapunov(mat, -np.array([[0,0],[0,B2 ]]) ) [1,1]
            demboth=solve_lyapunov(mat, -np.array([[B1,0],[0,B2 ]]) ) [1,1]
            envpred=solve_lyapunov(mat, -np.array([[0,0],[0,B2 ]])**2 ) [1,1]
            envboth=solve_lyapunov(mat, -np.array([[B1,0],[0,B2 ]])**2  ) [1,1]
            return dempred,demboth,envpred,envboth
        C22=np.array(zip(*[list(getinv(*args)) for args in zip(*[z.ravel() for z in (J11,J12,J21,J22,B1,B2) ]) ])).reshape((4,)+B1.shape)

        results=[T,D,np.real(eig),np.imag(eig)] +[np.clip(-np.log10(C22[z]**.5/B2),-0.5,None) for z in range(4) ]
        return results

    """======================== VECTOR FIELD ========================"""
    ngrid = 20
    prm={}
    prm.update(dft)
    idx=0
    plt.figure()
    for a,b in [[.3,.3],[.6,.6] ] :
        plt.subplot(1,2,1+idx)
        idx+=1
        prm['a']=a
        prm['b']=b
        prm['A']=(Amin(**prm)+Amax(**prm) )/2
        K1,K2 = geteq(**prm)[:2]
        u1,u2 = geteq(return_unstable=1,**prm)[:2]
        K1,K2=K1.squeeze(),K2.squeeze()
        X, Y = np.meshgrid(np.logspace(-2, np.log10(2*K1),ngrid), np.logspace(-2,np.log10( 20*K2),ngrid))
        dX,dY=eqs(np.log([X.ravel(),Y.ravel()]),**prm )
        norm=np.sqrt(dX**2+dY**2)
        dX,dY=dX/norm,dY/norm
        plt.quiver(X, Y, dX.reshape(X.shape), dY.reshape(Y.shape), color='b'),plt.xlabel(r'$B_1$'),plt.ylabel(r'$B_2$')
        plt.xscale('log')
        plt.yscale('log')
        plt.scatter(K1,K2,c='b')
        plt.scatter(u1,u2,c='r')
        plt.title(r'$\beta={}$, $\gamma={}$'.format(prm['a'],prm['b']) )


    """======================== T ========================"""
    ngrid = 100
    betas, gammas = np.meshgrid(np.linspace(0,1, ngrid)[1:-1],
                         np.linspace(0,1, ngrid)[1:-1] )

    prm={}
    prm.update(dft)
    B1,B2,err=geteq(a=betas,b=gammas,shape=betas.shape,**prm)
    stability=getstab(B1,B2,a=betas,b=gammas,**prm)
    LT, LD, Lreal, Limag=stability[:4]

    def pcolor(*args,**kwargs):
        kwargs.setdefault('rasterized',1)
        return plt.pcolormesh(*args,**kwargs)

    plt.subplot(341)
    plt.colorbar(pcolor(betas,gammas,np.log10( np.clip(B2,1,None) ) ))
    plt.gca().set_aspect(1)
    plt.xlabel(r'$\beta$'),plt.ylabel(r'$\gamma$')
    plt.subplot(345)
    vmax=np.max(np.abs(Lreal))
    plt.colorbar(pcolor(betas, gammas, Lreal,vmin=-vmax,vmax=vmax,cmap='seismic_r'))
    plt.gca().set_aspect(1)
    plt.xlabel(r'$\beta$'), plt.ylabel(r'$\gamma$'),plt.title("Real part")
    plt.subplot(346)
    vmax=np.max(np.abs(Limag))
    plt.colorbar(pcolor(betas, gammas, Limag,vmin=-vmax,vmax=vmax,cmap='seismic_r'))
    plt.gca().set_aspect(1)
    plt.xlabel(r'$\beta$'), plt.ylabel(r'$\gamma$'),plt.title("Imag part")
    plt.subplot(347)
    vmax=np.max(np.abs(LT))
    plt.colorbar(pcolor(betas,gammas,LT,vmin=-vmax,vmax=vmax,cmap='seismic_r'))
    plt.gca().set_aspect(1)
    plt.xlabel(r'$\beta$'), plt.ylabel(r'$\gamma$'),plt.title("Trace")
    plt.subplot(348)
    vmax=np.max(np.abs(LD))
    plt.colorbar(pcolor(betas,gammas,LD))
    plt.gca().set_aspect(1)
    plt.xlabel(r'$\beta$'), plt.ylabel(r'$\gamma$'),plt.title("Determinant")

    stabtitles=['Dem. pred.','Dem. both','Env. pred','Env. both' ]
    for z in range(4):
        plt.subplot(3,4,9+z)
        plt.colorbar(pcolor(betas,gammas,stability[4+z],linewidth=0))
        plt.gca().set_aspect(1)
        plt.xlabel(r'$\beta$'), plt.ylabel(r'$\gamma$'),plt.title(stabtitles[z])


    ngrid=5000
    prm={}
    prm.update(dft)
    prm['D1']=0.01
    plt.subplot(342)
    a=b=0.3
    prm['a']=a
    prm['b']=b
    amin,amax=Amin(  **prm), Amax( **prm)
    As=np.linspace(amin,3*amax, ngrid)
    prm['A']=As
    B1,B2,err= geteq(shape=As.shape,**prm )
    stab=getstab(B1,B2,**prm)[0]
    plt.plot(As[stab<0],np.log10(np.clip(B2[stab<0],1,None)))
    plt.plot(As, np.log10(np.clip(geteq(shape=As.shape,return_unstable=1 ,**prm ) [1],1,None)) ,linestyle='--')
    plt.title(r'Donor-limited ($\beta+\gamma<1$)'),plt.xlabel(r'Attack rate $\log_{10} \;A$'), plt.xticks([amin,amax],[r'$A_{\min}$',r'$A_{\max}$' ])
    plt.ylabel(r'$\log_{10} \; B_2$')
    
    plt.subplot(343)
    prm={}
    prm.update(dft)
    prm['D1']=0.01
    a=b=0.7
    prm['a']=a
    prm['b']=b
    amin,amax=Amin( **prm), Amax(  **prm)
    As=np.linspace(amin,2*amax, ngrid)
    prm['A']=As
    plt.plot(As,np.log10(np.clip(geteq(shape=As.shape,return_unstable=1 ,**prm )[1] ,1,None))  )
    plt.title(r'Predation-limited ($\beta+\gamma>1$)' ),plt.xlabel(r'Attack rate $\log_{10} \;A$'), plt.xticks([amin,amax],[r'$A_{\min}$',r'$A_{\max}$' ])
    plt.ylabel(r'$\log_{10}\; B_2$')

    plt.subplot(344)
    betas=np.linspace(0,1,ngrid)
    g1s=[0.5,1.,2.]
    bb=list(betas)*2
    for g1 in g1s:
        prm={}
        prm.update(dft)
        prm['g1']=g1
        plt.plot(bb,np.log10(np.concatenate([Amin(a=betas,b=0.5,**prm),Amax(a=betas,b=0.5,**prm) ]) ) )
        
    plt.legend([r'$g_1={}$'.format(g) for g in  g1s] ),plt.ylabel(r'Attack rate $\log_{10} \;A$'),plt.xlabel(r'$\beta$'),plt.title(r'Range $[A_{\min},A_{\max}]$')
    plt.show()


##### MODEL ANALYSIS
theo()

##### ADAPTIVE DYNAMICS
twopred(loop=True)
```
